# Roles of microbiota in autoimmunity in Arabidopsis

**DOI:** 10.1101/2023.03.06.531303

**Authors:** Yu Ti Cheng, Caitlin A. Thireault, Bradley C. Paasch, Li Zhang, Sheng Yang He

**Affiliations:** Department of Biology, Duke University, Durham, NC, USA; Howard Hughes Medical Institute, Duke University, Durham, NC, USA; Department of Energy Plant Research Laboratory, Michigan State University, East Lansing, MI, USA

**Author notes:** Corresponding authors; Yu Ti Cheng Sheng Yang He.

## Abstract

Over the past three decades, researchers have isolated plant mutants that display constitutively activated defense responses in the absence of pathogen infection. These mutants are called autoimmune mutants and are typically dwarf and/or bearing chlorotic/necrotic lesions. From a genetic screen for *Arabidopsis* genes involved in maintaining a normal leaf microbiota, we identified *TIP GROWTH DEFECTIVE 1* (*TIP1*), which encodes a S-acyltransferase, as a key player in guarding leaves against abnormal microbiota level and composition under high humidity conditions. The *tip1* mutant has several characteristic phenotypes of classical autoimmune mutants, including a dwarf stature, displaying lesions, and having a high basal level of defense gene expression. Gnotobiotic experiments revealed that the autoimmune phenotypes of the *tip1* mutant are largely dependent on the presence of microbiota as axenic *tip1* plants have markedly reduced autoimmune phenotypes. We found that the microbiota dependency of autoimmune phenotypes is shared by several “lesion mimic”-type autoimmune mutants in *Arabidopsis*. Interestingly, autoimmune phenotypes caused by mutations in *NLR* genes do not require the presence of microbiota and can even be partially alleviated by microbiota. Our results therefore suggest the existence of two classes of autoimmunity (microbiota-dependent vs. microbiota-independent) in plants. The observed interplay between autoimmunity and microbiota in the lesion mimic class of autoimmunity is reminiscent of the interactions between autoimmunity and dysbiosis in the animal kingdom.

## Main

In the past forty years, tremendous progress has been made in the understanding of plant immune responses against pathogens^1–3^. The plant innate immune system is comprised of both constitutive physical barriers and inducible immune responses. Inducible immunity can be initiated by plasma membrane-residing receptor kinases that recognize conserved microbe-associated molecular patterns, resulting in a broad-spectrum basal immunity called pattern-triggered immunity (PTI). Successful pathogens have evolved virulence-associated effector molecules to defeat the plant immune system and/or to create a conducive microenvironment within the host tissue as two major pathogenic mechanisms^4–7^. In response, plants have evolved an array of intracellular nucleotide-binding leucine-rich repeat (NLR) immune receptors that recognize the presence of specific pathogen effectors, leading to a more robust effector-triggered immunity (ETI) that often involved hypersensitive cell death^2,8^. Activation and mutual potentiation of PTI and ETI lead to accumulation of defense hormones, including salicylic acid, and activation of defense gene expression and other cellular responses^8,9^. Activation of plant immune response is often accompanied by growth inhibition, a phenomenon known as growth-defense trade-offs^10,11^

In nature, however, plants spend most of their life in an environment that is occupied by enormously diverse, mostly non-pathogenic (commensal) microorganisms^12,13^. Plant-associated commensal microbial community plays a vital role in influencing host growth, development, and stress responses^14,15^. However, compared to plant-pathogenic microbe interactions, less is known about how plants recognize and communicate with their surrounding non-pathogenic microbial communities and how plants fine tune their immune system to achieve a long-term, harmonious state in the context of complex microbial communities. Only in the past decade, thanks to the advent of high throughput sequencing technologies and availability of genetic mutants in model systems, such as *Arabidopsis* and rice, increasing efforts are being devoted to the study of the interplay between plant host genetics and associated microbial communities^12,14,16^. Nevertheless, the mechanisms of plant-microbiome interactions in terms of (i) how plants recognize, configure, and maintain a homeostatic composition of their associated microbiota and (ii) how the host immune system distinguishes non-self signals derived from commensal vs. pathogenic microbes are still largely unclear.

The lifestyle of commensal bacterial microbiota in plant leaves resembles those of non-pathogenic strains of phyllosphere bacterial phytopathogens: both are adapted to live in plant tissues but are unable to multiply to a high population level *in planta.* Recent studies have shown that the transcriptomes of commensal bacteria *in planta* share a high degree of similarity to that of a non-pathogenic mutant of the phyllosphere-adapted bacterial pathogen *Pseudomonas syringae* pv. *tomato* (*Pst*) DC3000^17,18^. Furthermore, two *Arabidopsis* quadruple mutants, *min7 fls2 efr cerk1* (*mfec*) and *min7 bak1 bkk1 cerk1* (*mbbc*), which are defective in pattern-triggered immunity and MIN7-associated intracellular vesicle trafficking, not only fail to control the proliferation of non-pathogenic mutants of *Pst* DC3000, but also unable to maintain a typical endophytic leaf microbiota^19,20^. In addition, immunity-associated reactive oxygen species (ROS) is required for maintaining a homeostatic leaf microbiota as the *Arabidopsis rbohD* mutant, which is defective in the generation of immunity-associated ROS, has altered leaf microbiota composition^21^. Taken together, these initial studies begin to identify plant genes/pathways that are required for maintaining a normal bacterial microbiota in *Arabidopsis* leaves and provide a strong link between plant immune regulation and microbiota homeostasis.

To identify additional plant genes involved in regulating plant-microbiome interactions and microbiota homeostasis, we conducted a forward genetic screen to isolate *Arabidopsis* mutants that exhibit an altered response to non-pathogenic mutants of *Pst* DC3000 and endophytic leaf microbiota. Characterization of the resulting mutants led to an unexpected broad connection between microbiota and autoimmunity in plants.

## Results

### Genetic screen to identify *Arabidopsis* mutants with altered leaf microbiota level

We previously found that *Arabidopsis* mutant plants compromised in three pattern-recognition co-receptor genes, *BAK1*, *BKK1* and *CERK1*, are still capable of preventing over-proliferation of non-pathogenic strain *Pst* D28E^19^ and endogenous leaf microbiota^20^. We therefore conducted a forward genetic screen in the *bak1-5 bkk1-1 cerk1-2* triple mutant background^22,23^ (*bbc* hereafter), intending to identify genes whose functions are either independent of and/or additive to these three pattern-recognition co-receptors (see Extended Data Figure 1 for the setup of the genetic screen).

The primary screen was carried out by flood-inoculating 3-week-old plate-grown M2 seedlings with *Pst* D28E, a non-pathogenic mutant which was constructed by deleting 28 effectors of *Pst* DC3000^24^. Individual plants that showed disease-like symptoms (e.g., necrosis or chlorosis) were transplanted to soil and grown for seeds. Secondary screen of putative mutants was conducted by monitoring disease-like symptoms after syringe-infiltration of *Pst* D28E into leaves of 4-week-old soil-grown M3 plants. We also monitored disease-like symptoms induced by natural soil-derived microbiota under holoxenic conditions in a peat-based gnotobiotic system^25^. In total, we identified ten mutants with various degrees of enhanced disease-like symptoms in response to non-pathogenic *Pst* D28E and/or natural soil-derived microbiota; we named them *<u>g</u>uardian of no<u>r</u>mal <u>m</u>icrobiota* (*grm*) mutants.

### Characterization of the *grm1* mutant

Next, we conducted detailed characterization of one of the identified *grm* mutants, *grm1*. When grown in soil, the *grm1* mutant had a smaller stature compared to its progenitor, the *bbc* triple mutant. Notably, lower leaves that were in contact or in close proximity to the soil showed mild chlorosis (Fig. 1a, top panel). Previously, we found that *mfec* and *mbbc* mutant plants exhibited dysbiotic endophytic leaf microbiota and leaf tissue damage when grown under high humidity, a common environmental condition associated with plant disease outbreaks in nature^19,20^. The chlorotic lower leaves of the *grm1* mutant resembled those of *mfec* and *mbbc* mutant plants and prompted us to investigate if *grm1* also displays dysbiotic endophytic leaf microbiota under high humidity. As expected in wild type *Arabidopsis* (accession Col-0, which is the progenitor of the *bbc* triple mutant), characteristic hyponastic changes in leaf morphology were induced after five days of high humidity treatment (~95% relative humidity [RH]; Fig. 1a, lower panel); however, high humidity was not able to cause over-proliferation of endophytic leaf microbiota and no leaf chlorosis nor necrosis was observed (Fig. 1b). Similar to Col-0 plants, the *bbc* mutant also maintained a low level of culturable endophytic leaf microbiota, similar to these plants under ambient humidity (~50% RH). In contrast, after five days of high humidity treatment, most of the *grm1* true leaves showed strong chlorosis (Fig. 1a, lower panel). Quantification of culturable endophytic microbiota loads by colony counts on agar plates revealed that, compared to Col-0 and the *bbc* mutant, the endophytic microbiota within leaves of *grm1* plants grown under high humidity was more than three orders of magnitude higher (Fig. 1b). In addition to a drastic increase in leaf endophytic microbiota, the relative abundance of leaf microbiota members in the *grm1* mutant also shifted overwhelmingly to Proteobacteria (Fig. 1c, ~97% in *grm1* leaves compared to ~45-55% in Col-0 and *bbc* leaves). In the *grm1* mutant, amplicon sequence variants (ASVs) in Proteobacteria belong predominantly to the genus *Pseudomonas* of the class Gammaproteobacteria, while *Bacillus* and *Paenibacillus* belonging to Firmicutes became nearly undetectable (Fig. 1c and Extended Data Table 1). Shannon index that measures richness and evenness of a microbial community composition also decreased in the *grm1* mutant, reflecting the overwhelming presence of Proteobacteria (Fig. 1d).

**Figure 1.**
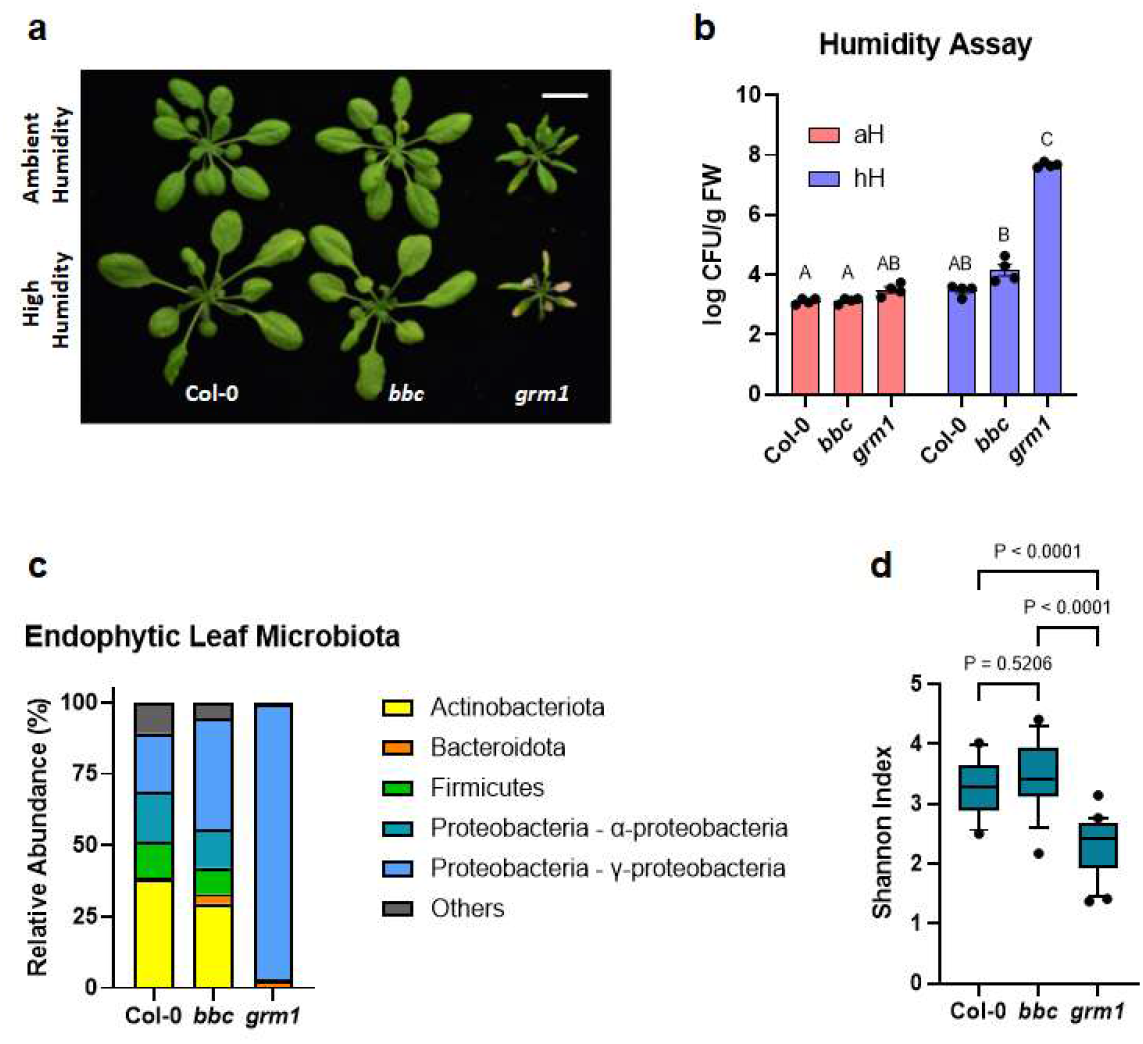
The appearance and leaf microbiota phenotypes of the *grm1* mutant. **a,** Top panel, four-week-old soil-grown Col-0, *bbc* and *grm1* plants under ambient humidity (~50% RH) for five days (basal condition, controls). Bottom panel, four-week-old soil-grown plants shifted to high humidity (~95% RH) for five days. Images were taken on day five of the treatments. Scale bar equals 2 cm. **b,** Population sizes of endophytic leaf microbiota after five days of indicated humidity conditions. aH = ambient humidity (~50% RH; basal condition, controls); hH = high humidity (~95% RH). Each column represents bacterial titer as log-transformed colony forming units (CFU) per gram of fresh weight (FW). Data are displayed as mean ± SEM (n=4 biological replicates; each biological replicate contains 1-3 leaves from one plant). Different letters represent significant differences (p < 0.05, two-way ANOVA with Tukey’s HSD test). Experiment was independently performed three times with similar results. **c,** Relative abundance of endophytic leaf bacteria at the phylum level and at the class level for Proteobacteria. **d,** Shannon indexes of endophytic leaf bacteria based on 16S rDNA amplicon sequence profiling of indicated genotypes. The center lines of the box plot represent means, the box edges are the 75^th^ and 25^th^ percentiles, whiskers extend to 10-90 percentiles, and dots are outliers.

### Identification of the causal mutation in the *grm1* mutant

To identify the causative mutation in the *grm1* mutant, *grm1* plants were backcrossed with its progenitor, the *bbc* triple mutant, to generate a segregated F2 population. Analyzing the mapping-by-sequencing data from *grm1* co-segregates revealed that the mutation was located on chromosome 5 between 5Mb and 8Mb with the allele frequency of the *grm1*-like pool peaking around 7Mb (Extended Data Fig. 2a; see Extended Data Table 2 for a list of mutated loci in this region). Of all candidates, a G to A mutation on chromosome 5 at position 6,877,509 has the strongest effect. This G to A mutation occurs at the splicing junction of the 5’ end of the third intron of the *TIP GROWTH DEFECTIVE 1* (*TIP1*) gene and is expected to disrupt the splicing pattern, leading to a premature stop codon instead of tryptophan at position 171 (Extended Data Fig. 4a). RT-PCR using primers flanking the mutation locus followed by Sanger sequencing revealed that, indeed, the mutation altered the splicing pattern of the *TIP1* transcript (Extended Data Fig. 2b). Transgene complementation with the full length *TIP1* gene driven by its native promoter could complement *grm1* mutant phenotypes (i.e., reversion of the dwarf stature and humidity-dependent dysbiotic phenotypes to those in *bbc* and wild-type Col-0), confirming the causative mutation in *grm1* is in the *TIP1* gene (Extended Data Fig. 3).

To determine if the dysbiotic phenotypes of the *grm1* mutant (i.e., the *bbc tip1* quadruple mutant) are dependent on the background *bbc* mutations, we segregated the *tip1* mutation from the *bbc* triple mutations by outcrossing the *grm1* mutant with wild-type Col-0 plants and genotyping the resulting F2 population (see Extended Data Table 5 for primes used to genotyping). From The Arabidopsis Biological Resource Center (ABRC) ^26^, we also obtained two independent *tip1* single mutant alleles (SALK_020996 and SALK_052842) carrying T-DNA insertions in the *TIP1* gene (Fig. 2a). All three *tip1* single mutants were larger than the original *grm1* (*bbc tip1*) mutant, but still smaller than wild type Col-0 plants (Fig. 2b). Interestingly, the humidity-dependent dysbiosis phenotypes (e.g., leaf chlorosis) (Extended Data Fig. 4b) and over-proliferation of endophytic leaf microbiota were still observed in all three alleles of *tip1* single mutant plants, suggesting dysbiotic phenotypes observed in *grm1* do not require *bbc* triple mutations in the background (Fig. 2c). Since all three mutant alleles of *tip1* behaved similarly, we used *tip1*^W171STOP^, the single mutant allele isolated from this screen, for all subsequent experiments.

**Figure 2.**
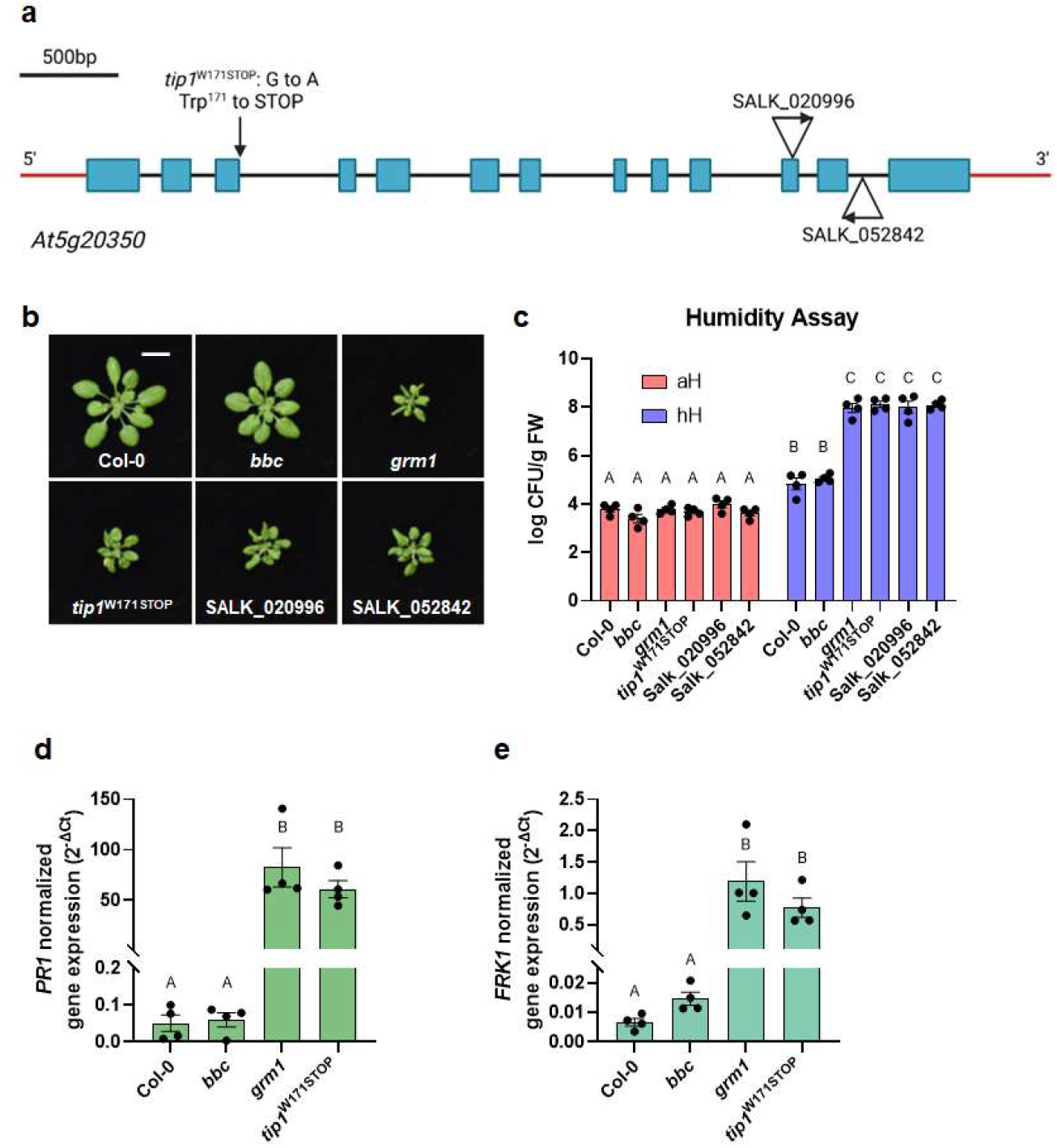
Characterization of *tip1* single mutant plants. **a,** A schematic diagram showing various mutant alleles in the *TIP1* gene. *tip1*^W171STOP^ is the allele isolated from this study which contains a G to A mutation at the splicing junction that is expected to cause a pre-mature STOP codon at amino acid residue Trp^171^ in the ankyrin-repeat domain. SALK_020996 and SALK_052842 are T-DNA insertion alleles obtained from ABRC. Created with BioRender.com. **b,** Images of four-week-old, soil-grown Col-0, *bbc*, *grm1* and *tip1* single mutant plants. Scale bar equals 2 cm. **c,** Population sizes of endophytic leaf microbiota after five days under humidity conditions indicated. aH = ambient humidity (~50% RH; basal condition, controls); hH = high humidity (~95% RH). Results represent the mean values ± SEM (n=4 biological replicates; each biological replicate contains 1-3 leaves from one plant). Different letters represent a significant difference (p < 0.05, two-way ANOVA with Tukey’s HSD test). Experiment was independently performed three times with similar results. **d,e,** Expression levels of *PR1* (d) and *FRK1* (e) genes in four-week-old, soil-grown Col-0, *bbc*, *grm1* and *tip1* plants. *PP2AA3* expression was used for normalization. Results represent the mean values ± SEM of four biological replicates. Each biological replicate is a pool of three plants. Different letters represent a significant difference (p < 0.05, one-way ANOVA with Tukey’s HSD test). Experiment was independently performed twice with similar results.

### The *tip1* mutant has features of autoimmune mutants

We noticed that the morphological phenotypes of *grm1* and *tip1* (i.e., small statures and chlorotic leaves) were reminiscent of typical autoimmune mutants, which have been isolated in the past few decades. One hallmark of autoimmunity is constitutive high basal expression of immune-related marker genes in the absence of pathogen attacks. We therefore analyzed two immune-related molecular markers, *Pathogenesis-Related Gene 1* (*PR1*) and *flg22-Induced Receptor-like Kinase 1* (*FRK1*). Indeed, both *grm1* and *tip1* plants have heightened *PR1* and *FRK1* expression under basal condition (i.e., in the absence of pathogen inoculation) (Fig. 2d and 2e). *TIP1* gene expression itself was found to be induced by flg22, a flagellin-derived peptide that induces many PTI-associated genes^27,28^ (Extended Data Fig. 5).

### *tip1*-mediated autoimmunity is distinct from *snc1*-mediated autoimmunity

We were intrigued by the morphological phenotypes and heightened immune-related marker gene expression of the *tip1* mutant as they point to a connection between dysbiosis and autoimmunity. However, it is not known (i) if all autoimmune mutants possess a defect in maintaining microbiota homeostasis and/or (ii) if their autoimmune phenotype is dependent on microbiota. To investigate on a possible connection between altered microbiome and autoimmunity, we first examined microbiota-related phenotypes of a widely studied *Arabidopsis* autoimmune mutant, *snc1*, which contains a gain-of-function E552K mutation that results in an elevated level of the SNC1^E552K^ protein^29,30^.

As expected, *snc1* mutant plants had a dwarf morphology (Fig. 4a, top panel), constitutively elevated immune-related marker gene expression (Fig. 3a and 3b) and enhanced resistance to the virulent pathogen *Pst* DC3000 (Fig. 3c). However, *snc1* and *tip1* mutant plants behaved differently in response to non-pathogenic bacteria. As shown in Figure 3d, *tip1* single mutant plants were more susceptible to the non-pathogenic *Pst* Δ*hrcC* mutant strain (*Pst* DC3000 defective in type III secretion system^31^), whereas *snc1* plants were marginally more resistant to the *Pst* Δ*hrcC* mutant compared to wild-type Col-0 plants.

**Figure 3.**
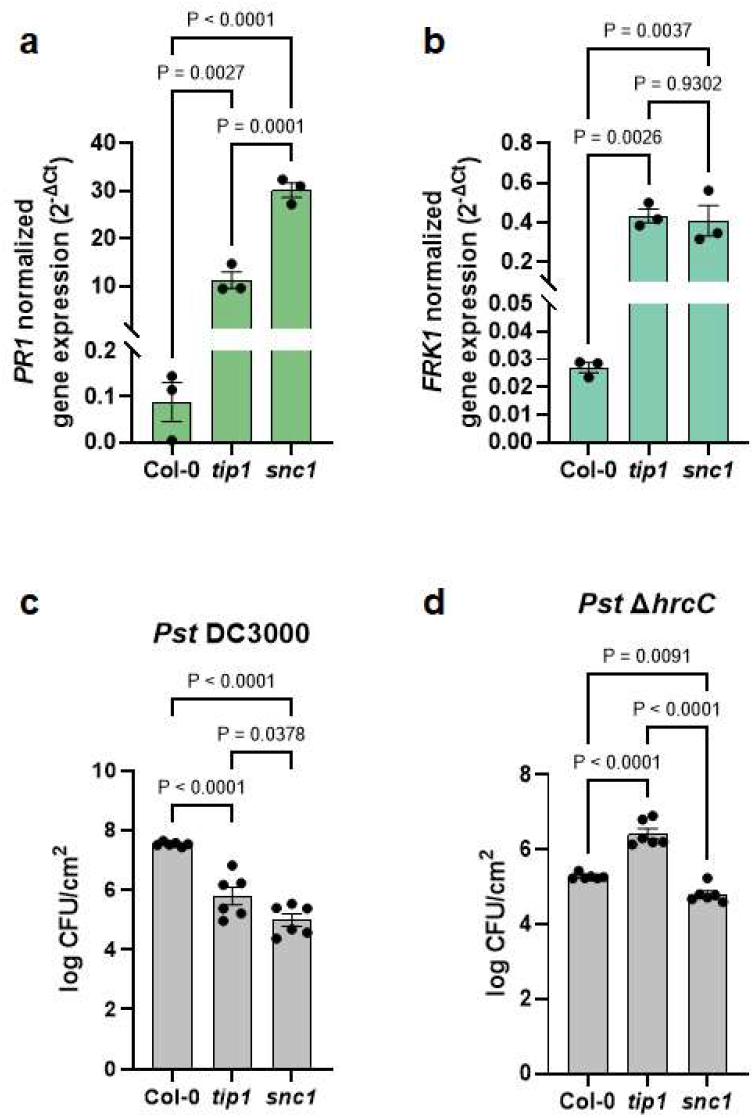
The autoimmune phenotypes of *tip1* and *snc1* mutants. **a,b,** Expression level of *PR1* (a) and *FRK1* (b) genes in four-week-old, soil-grown Col-0, *tip1* and *snc1* plants. *PP2AA3* was used for normalization. Results represent the mean values ± SEM of three biological replicates. Each biological replicate is a pool of three plants. Statistical analysis was performed using one-way ANOVA with Tukey’s HSD test. Experiment was independently performed twice with similar results. **c,d,** Total bacterial populations in Col-0, *tip1* and *snc1* leaves three days after *Pst* DC3000 (c) or five days after *Pst* Δ*hrcC* (d) infiltration. Humidity was kept at ~95% throughout the duration of the disease assay. Each column represents bacterial titer as log-transformed colony forming units (CFU) per cm^2^ and is the mean of six biological replicates; each biological replicate contains leaf discs from infiltrated leaves from one plant; total of six plants were infiltrated. Error bars indicate SEM. Statistical analysis was performed using one-way ANOVA with Tukey’s HSD test. Experiment was independently performed three times with similar results.

**Figure 4.**
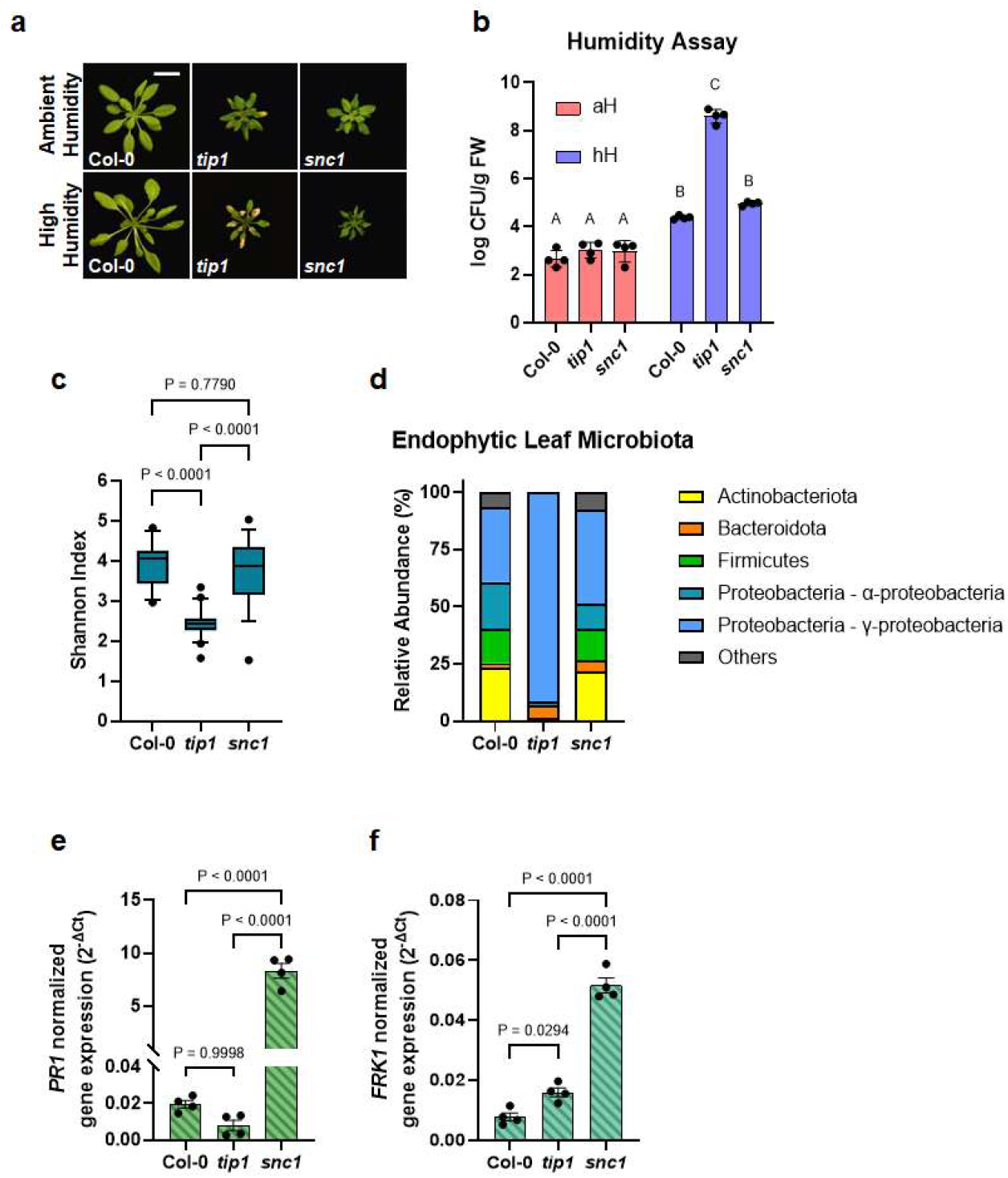
The distinct autoimmune phenotypes of *tip1* and *snc1* mutants. **a,** Top panel, four-week-old, soil-grown Col-0, *tip1* and *snc1* plants under ambient humidity (~50% RH) for five days (basal condition, controls). Bottom panel, four-week-old, soil-grown plants shifted to high humidity (~95% RH) for five days. Images were taken on day five of the treatments. Scale bar equals 2 cm. **b,** Population sizes of endophytic leaf microbiota after five days of plant growth under humidity conditions indicated. aH = ambient humidity (~50% RH; basal condition, controls); hH = high humidity (~95% RH). Results represent the mean values ± SEM of four biological replicates; each biological replicate contains 1-2 leaves from one plant. Different letters represent a significant difference (p < 0.05, two-way ANOVA with Tukey’s HSD test). Experiment was independently performed three times with similar results. **c,d,** Shannon indexes (c) and relative abundance (d) of endophytic bacterial microbiota at the phylum level and at class level for Proteobacteria of in Col-0, *tip1* and *snc1* leaves based on 16S rDNA amplicon sequence profiling. The center lines of the box plot represent means, the box edges are the 75^th^ and 25^th^ percentiles, whiskers extend to 10-90 percentiles, and dots are outliers. **e,f,** Expression level of *PR1* (e) and *FRK1* (f) genes in 2.5-week-old plate-grown Col-0, *tip1* and *snc1* plants. *PP2AA3* expression was used for normalization. Results represent the mean values ± SEM of four biological replicates. Each biological replicate is a pool of two seedlings. Statistical analysis by one-way ANOVA with Tukey’s HSD test. Experiment was independently performed twice with similar results.

Furthermore, while most *tip1* leaves showed severe chlorotic lesions when shifted to high humidity condition for five days, *snc1* mutant plants were morphologically similar under either ambient or high humidity conditions (Fig. 4a). In addition, enumeration of leaf endophytic microbiota showed that *snc1* plants carried similar levels of culturable bacteria inside their leaves as that of Col-0 plants, whereas *tip1* plants had more than 1,000-fold increase of endophytic bacterial load compared to Col-0 after higher humidity shift (Fig. 4b). The differences in leaf endophytic microbiota between *tip1* and *snc1* plants were not just in quantity but also in composition. Profiling bacterial communities inside Col-0, *tip1* and *snc1* leaves using 16S rDNA amplicon sequencing revealed that, compared to Col-0, *tip1* plants had substantially reduced leaf endophytic microbiota diversity (Fig. 4c) with overwhelming relative abundance of Gammaproteobacteria (Fig. 4d and Extended Data Table 3). In contrast, *snc1* plants had a diverse leaf endophytic microbiota composition, similar to that of Col-0 plants (Fig. 4c and 4d). Finally, when grown in aseptic agar plates, *snc1* plants continued to exhibit heightened immune-related marker gene expression in the absence of microbes, whereas *PR1* and *FRK1* expression in *tip1* mutant plants greatly subsided to close to the low levels observed in wild type Col-0 plants (Fig. 4e and 4f).

### A broad role of microbiota in “lesion-mimic” autoimmunity in *Arabidopsis*

The interesting contrast in microbiota-dependency for autoimmune phenotypes between the *tip1* vs. *snc1* mutants prompted us to further investigate if there is a broad connection between microbiota-dependency and autoimmunity in other reported *Arabidopsis* autoimmune mutants^32,33^ (see Extended Data Table 4 for list of mutants and stock numbers). Based on if they possessed tissue lesions when grew in potting soil under standard growth chamber conditions, we found that the *tip1*-like autoimmune mutant category includes (i) the *aca4 aca11* double mutant, which harbors mutations on two vacuolar calcium ion pumps ACA4 and ACA11^34^, (ii) the *acd5* mutant, which carries mutation in a ceramide kinase^35,36^ and (iii) the *lsd1* mutant, which has a defective zinc-finger protein in *Arabidopsis*^37^. The *snc1*-like autoimmune mutant category consists of (i) the *chs3* mutant, which carries a gain-of-function mutation in a TIR-type NLR immune receptor^38^, (ii) *dnd1*^39^ and (iii) *dnd2*^40^ mutants, which carry mutations in two cyclic nucleotide-gated cation channels. Mutant plants in the *tip1*-like category showed severe leaf lesions (Fig. 5a; top panel) and harboured high levels of endophytic leaf microbiota under high humidity (Fig. 5b), whereas mutant plants in the *snc1*-like category had no visible lesions (Fig. 5a; bottom panel) and carried low levels of endophytic leaf microbiota similar to wild-type Col-0 plants (Fig. 5c).

**Figure 5.**
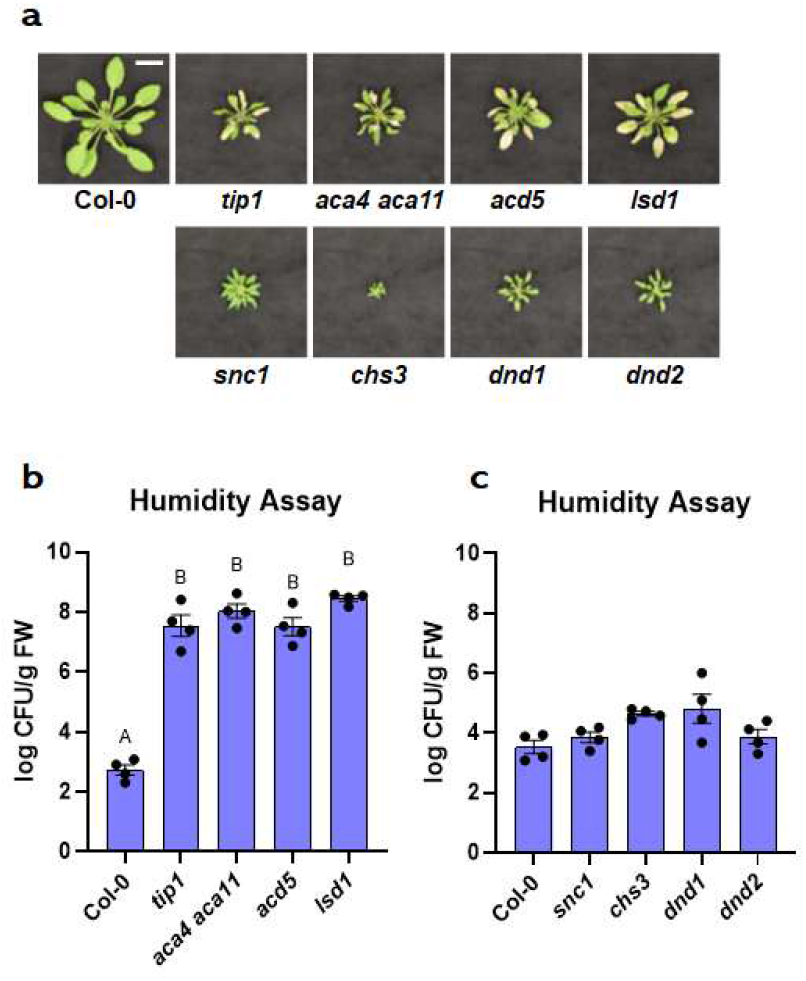
The appearance and leaf microbiota phenotypes of *Arabidopsis* autoimmune mutants. **a,** Images of four-week-old, soil-grown *Arabidopsis* autoimmune mutants exposed to high humidity (~95% RH) for five days. Top panel, Col-0, *tip1* and three previously identified “lesion-mimic” autoimmune mutants; bottom panel, *snc1* and three previously identified autoimmune mutants that showed no visible lesions. Scale bar equals 2 cm. **b,** Population sizes of endophytic leaf microbiota after five days of plant growth under high humidity condition (~95% RH) in *tip1* and three previously identified “lesion-mimic” autoimmune mutants. **c,** Population sizes of endophytic leaf microbiota after five days of plant growth under high humidity in *snc1* and three previously identified autoimmune mutants with no visible lesions. Results represent the mean values ± SEM of four biological replicates; each biological replicate contains 1-3 leaves from one plant. Different letters represent a significant difference (p<0.05, one-way ANOVA with Tukey’s HSD test). Experiment was independently performed three times with similar results.

To further characterize a possible microbiota dependency of autoimmune phenotypes in these two categories of mutants, we grew mutant plants in the absence (axenic) or presence (holoxenic) of a natural soil-derived microbiota using GnotoPots, a peat-based gnotobiotic system as recently described^25^. Growing under the holoxenic condition, lesion-mimic mutants showed various degrees of chlorosis and lesions (Fig. 6a, top panel, and Extended Data Fig. 7a) and heightened immune-related marker gene expression (Fig. 6c; right panel). However, in the absence of microbiota, these autoimmune mutants showed neither chlorosis nor lesions (Fig. 6a, bottom panel) and their immune marker gene expression also subsided to a low basal level, with exception of *lsd1*(Fig. 6c, left panel). In contrast, mutants in the *snc1*-like category have high basal *PR1* expression even in the axenic condition. For example, the *chs3* mutant showed heightened *PR1* expression regardless of presence or absence of microbiota (Fig. 6d), behaving similarly to the *snc1* mutant which shows microbiota-independent autoimmunity. Furthermore, compared to microbiota-induced lesions and immune-related gene expression in lesion mimic mutants, the autoimmune dwarf phenotype of *chs3* mutants was noticeably alleviated in the presence of microbiota (Fig. 6b), again similar to the *snc1* mutant. In *dnd1* and *dnd2* mutants, *PR1* expression was elevated to a higher level when they were grown in the presence of microbiota compared to when grown in the axenic condition (Fig. 6d), behaving intermediately between lesion mimic-type and *snc1*-type. Together, these results suggest that there are at least two types of autoimmunity in plants, one depends on microbiota for autoimmune phenotypes, as exemplified by the *tip1* mutant, and the other is independent of microbiota, as exemplified by the *snc1* mutant. *dnd1* and *dnd2* mutants share with *snc1*-type in that they do not show lesions in the presence of microbiota and have a high basal defense gene expression in the absence of microbiota, although defense gene expression is further enhanced in the presence of microbiota.

**Figure 6.**
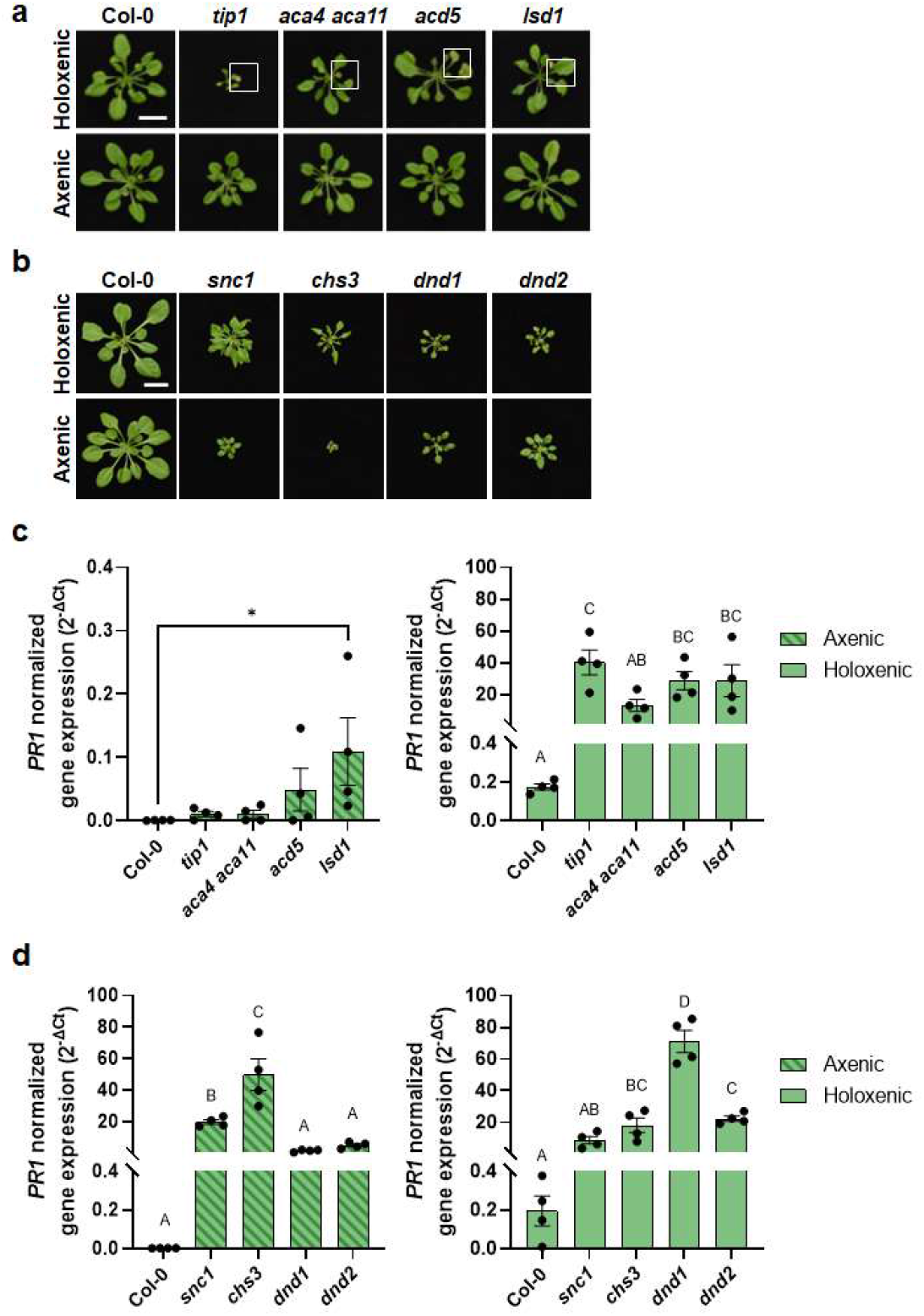
Microbiota dependency for autoimmunity in *Arabidopsis* autoimmune mutants. **a,** Five-week-old Col-0, *tip1* and three previously identified lesion-mimic autoimmune mutants grown in GnotoPots under holoxenic (top panel) or axenic (lower panel) conditions. Scale bar equals 2 cm. Zoomed in images (white squares) on leaf lesions are shown in Extended Data Figure 7a. **b,** Five-week-old Col-0, *snc1* and three previously identified autoimmune mutants that showed no visible lesions were grown in GnotoPots under holoxenic (upper panel) or axenic (lower panel) conditions. Scale bar equals 2 cm. **c,d,** *PR1* expression in *tip1* and three previously identified lesion-mimic autoimmune mutants (c) and *snc1* and three autoimmune mutants (d) grown in GnotoPots under axenic (left; with diagonal stripe pattern) or holoxenic (right) conditions. Results represent the mean values ± SEM of four biological replicates. Each biological replicate is a pool of two plants. Different letters and “*” represent a significant difference (p<0.05, one-way ANOVA). Experiment was independently performed twice with similar results.

### Microbiota dependency of autoimmunity in natural *Arabidopsis* accessions

Autoimmunity has been observed in natural *Arabidopsis* populations/accessions^41^. For example, *A. thaliana* accessions Est-1 and C24 have constitutively elevated defense gene expression and enhanced disease resistance toward the virulent pathogen *Pst* DC3000 when grown in potting soil^42,43^. We were therefore interested in knowing if autoimmune phenotypes of natural accessions are dependent on microbiota. When grown in potting soil under ambient humidity, Est-1 showed chlorosis and lesions on older leaves, whereas C24 had curly leaves and small stature but did not show chlorosis or lesions (Extended Data Fig. 6). However, like the *tip1* mutant, Est-1 leaves showed stronger leaf lesions under high humidity (Fig. 7a) and harboured a higher level of endophytic bacterial microbiota compared to Col-0 plants (Fig. 7b). In contrast, C24 plants did not show chlorosis or necrosis under ambient (Extended Data Fig. 6) or high humidity (Fig. 7a); additionally, similarly to the *snc1* mutant, C24 plants maintained similar levels of endophytic leaf bacterial microbiota compared to Col-0 (Fig. 7b). The phenotypic resemblance between *tip1* and Est-1 and between *snc1* and C24 prompted us to investigate if microbiota is required for the autoimmune phenotypes in Est-1 and C24. As shown in Figure 7c, under holoxenic condition, Est-1 shows lesions on leaves, albeit to a lesser extent compared to Est-1 grown under the conventional potting soil growth condition (Extended Data Fig. 6 and Extended Data Fig. 7b). Interestingly, like *tip1*, Est-1 plants did not show leaf lesions under the axenic condition. Furthermore, like the *tip1* mutant, the heightened *PR1* expression in Est-1 subsided to a low level when grown in the axenic condition (Fig. 7d; left panel). Conversely, like the *snc1* mutant, C24 plants had elevated *PR1* expression regardless of growth in the presence or absence of a microbial community (Fig. 7d). Another similarity between the *snc1* mutant and C24, which is in contrast to the *tip1* mutant, is the alleviation of their stunted growth morphology in the presence of microbiota (Fig. 6b and 7c).

**Figure 7.**
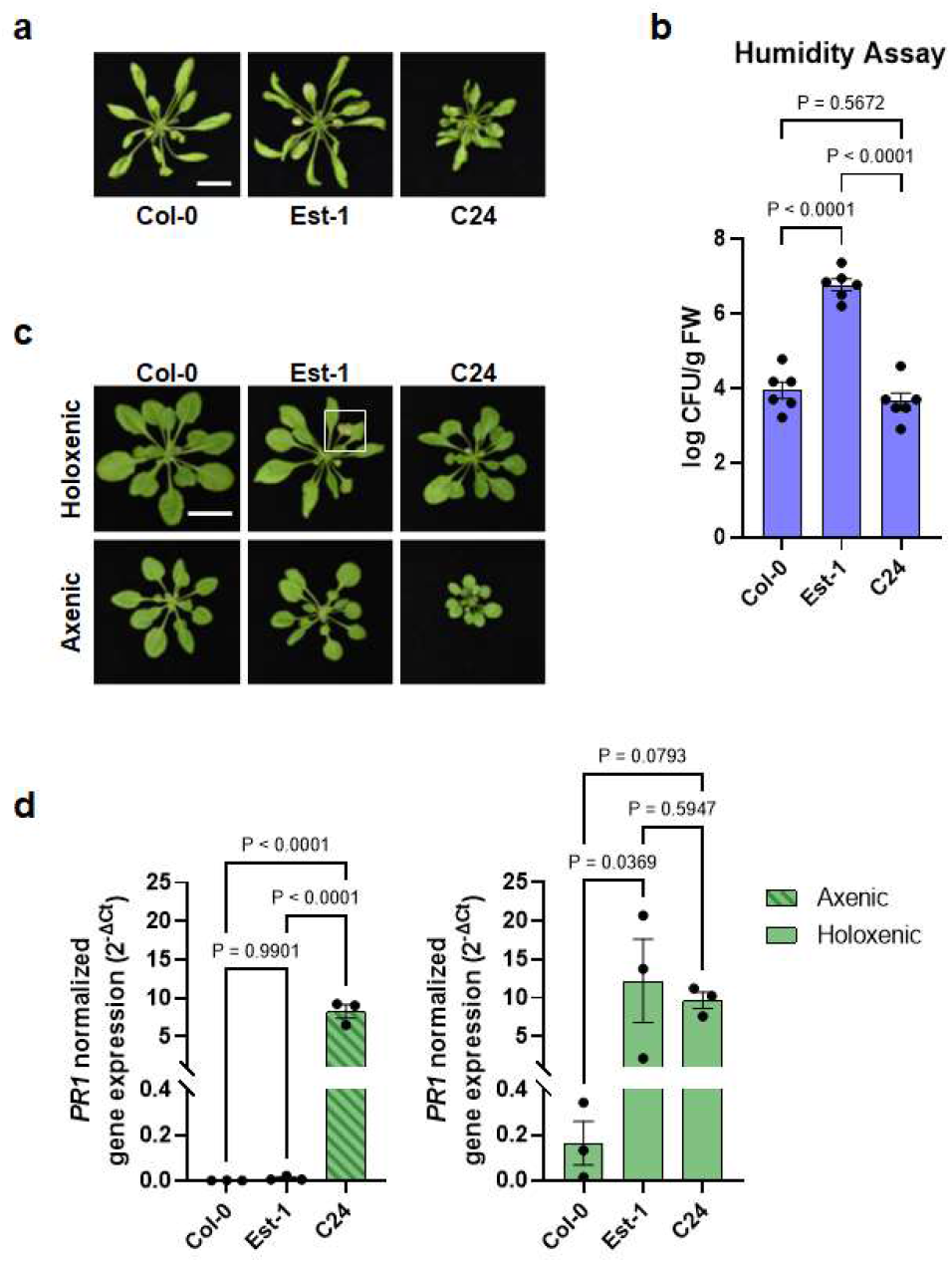
Microbiota dependency for autoimmunity in *Arabidopsis* natural accessions. **a,** Images of five-week-old, potting soil-grown *Arabidopsis* accessions (Col-0, Est-1 and C24) exposed to high humidity (~95% RH) for seven days. Scale bar equals 2 cm. **b,** Population sizes of leaf endophytic microbiota after seven days of plant growth under high humidity condition (~95% RH). Results represent the mean values ± SEM of six plants. Statistical analysis was performed using one-way ANOVA with Tukey’s HSD test. Experiment was independently performed three times with similar results. **c,** Five-week-old *Arabidopsis* accessions grown using GnotoPots under holoxenic (upper panel) or axenic (lower panel) conditions. Scale bar equals 2 cm. Zoomed in image (white square) on Est-1 leaf lesions is shown in Extended Data Figure 7b. **d,** *PR1* expression in *Arabidopsis* accessions grown in GnotoPots under axenic (left; with diagonal stripe pattern) or holoxenic (right) conditions. Results represent the mean values ± SEM of three biological replicates. Each biological replicate is a pool of two plants. Statistical analysis by one-way ANOVA with Tukey’s HSD test. Experiment was independently performed twice with similar results.

## Discussion

A healthy microbiome can play a vital role in initiating, training, and maintaining host immune homeostasis. In return, the host immune system can fine tune its immune strength to accommodate commensal/symbiotic microbes and to prepare for a robust immune response against pathogenic microbe invasion. As plants spend most of their life interacting with a vast number of commensal microbes and occasionally encountering pathogens, understanding the intricate interplays between plant immunity and the endophytic commensal microbiota is important for explaining how plants dial their plant immune system to maximize the effectiveness of plant immune responses to nurture beneficial microbes and/or fight against pathogens. In this study, we conducted a forward genetic screen aimed at identifying *Arabidopsis* mutants that cannot maintain a normal leaf microbiota. Among putative mutants isolated was *grm1*, which we characterized in detail.

The *grm1* mutant contains a missense mutation in the *TIP1* gene that encodes a S-acyltransferase. The first mutant allele of *TIP1* was isolated in a genetic screen for mutants that had defects in root hair development^44^. The causal mutation was later mapped to *At5g20350*^45^, which encodes one of the 23 DHHC-containing S-acyltransferases and is the only ankyrin-repeat containing DHHC S-acyltransferase in *Arabidopsis*. Both the *Arabidopsis* TIP1 protein and human HIP14 (huntingtin-interacting protein 14; zDHHC17) have been shown to be functional orthologs of yeast Akr1p^45,46^. Akr1p and zDHHC17 are involved in vesicle trafficking. For example, acylation of yeast Yck2 protein by Akr1p is required for proper localization of Yck2 to the plasma membrane via secretory vesicles and Yck2p’s membrane association is essential for its biological function in yeast morphogenesis^47^.

Prior to this study, however, the connection between TIP1 and leaf microbiota homeostasis was not known. A comprehensive *Arabidopsis* acylome using multiple tissue types identified close to 1,100 putative S-acylated proteins^48^. Thirty seven percent of identified proteins overlapped with those identified in a previous study which used *Arabidopsis* root cell culture^49^. Of note, many of the identified proteins have been demonstrated to be associated with microbe perception and plant immune responses^49^. However, it is not known if S-acylation is required for their function and, so far, none of them are confirmed to be TIP1-specific substrates. In light of our findings on a genetic connection between *TIP1* and maintenance of a normal leaf microbiota, future research is needed to identify specific TIP1 substrate(s) that is required for microbiota homeostasis in *Arabidopsis* leaves.

A key finding in this study is that not only the *tip* mutants are unable to control the proliferation or maintain a normal composition of a leaf microbiota, but they also display dysbiosis-associated tissue damages and autoimmunity in the presence of microbiota (Fig. 4). The microbiota-dependent autoimmune phenotypes of the *tip1* mutant led us to broadly examine a potential connection between microbiota and previously reported “autoimmune” mutants in *Arabidopsis*. Based on how they respond to the existence of microbiota, it appears that autoimmune mutants in *Arabidopsis* can be divided into at least two classes (Extended Data Fig. 8). One class, exemplified by the *tip1* mutant, exhibits microbiome-dependent autoimmunity. The autoimmune phenotypes in this class largely disappeared when grown in the axenic conditions. Given that these mutants also have an increased microbiota load, this result suggests that the autoimmune phenotypes in this class of mutants are a consequence of harbouring an overabundant microbial community. The other class of autoimmunity is independent of microbiota and is represented by the *snc1* mutant. The autoimmune phenotypes of this class do not require the presence of microbiota. i.e., they have small statures and high *PR1* expression regardless of presence or absence of microbial communities. In fact, the presence of microbiota alleviates the stunted growth morphology of *snc1* and *chs3* (Fig. 6b), which is in striking contrast to those of the *tip1* class (Fig. 6a).

We find it interesting that microbiota-dependent and -independent autoimmunity can also be observed in *Arabidopsis* natural accessions. Although autoimmunity is often associated with fitness trade-offs at the individual plant level^10,11^, the presence of heightened basal immunity in natural accessions suggests a fitness advantage at the population level^43^. For example, if a devastating disease spreads through a largely susceptible *Arabidopsis* population, accessions like C24 and Est-1 may be able to survive and reproduce to avoid extinction of the entire population. The microbiota-dependent and -independent expression of autoimmunity in Est-1 vs. C24, as observed in our study (Fig. 7), may reflect different paths by which the two types of autoimmunity have convergently evolved in natural populations under different abiotic and biotic pressures. Future research may uncover other natural accessions that show a continuum range of autoimmune phenotypes in terms of microbiota-dependency, as observed for the *dnd1* and *dnd2* mutants, which exhibit microbiota-amplification of heighted basal defense gene expression (Fig. 6d).

Overall, results from this study begin to illustrate conceptual parallels in microbiota interactions with plants and animals and may have broad implications in understanding host-microbiota interactions in general. In mammalian-microbiota interactions, for example, dysbiotic shifting/reducing diversity in microbiome composition is often associated with inflammation and dysregulated immune responses that fail to distinguish self from non-self, which are characteristics of autoimmune disorders^50–52^. The similarities in autoimmune symptoms between *Arabidopsis tip1* and mammalian inflammatory autoimmunity are notable that they include a dysbiotic microbial community, tissue lesions, and dysregulated immune responses. The most renowned substrate of zDHHC17, the human functional homologue of *Arabidopsis* TIP1, is huntingtin (htt) ^46,53^. In light of the connection between *tip1* mutation and autoimmunity in *Arabidopsis*, it would be interesting for future research to examine if zDHHC17 mutations are associated with dysbiosis and/or if dysbiosis is involved in Huntington’s disease development. The advance in biochemistry studies of zDHHC17^54–56^ and the more easily amenable mutant studies at the whole organism level in *Arabidopsis* should facilitate further understanding of a possibly broad role of DHHC S-acyltransferases in microbiome homeostasis and immunity across the kingdoms of life.

## Methods

### Plant materials and growth condition

All seeds were surface-sterilized using 15% diluted bleach (containing final concentration of 1.2% active ingredient sodium hypochlorite [NaOCl]) before being sown onto potting soil. All plants were grown under a 12h day/12h night regimen with 100 µE light intensity and ~50% relative humidity, unless otherwise indicated. See Extended Data Table 4 for a complete list of *Arabidopsis* mutants and accessions used in this study.

For peat-based gnotobiotic experiments, plants were grown in GnotoPots^25^. Nutrients were supplemented with buffered half-strength Linsmaier and Skoog liquid media (0.5x LS; Caisson Labs LSP03). Soil for natural microbiota inoculation was harvested from a miscanthus field plot at Michigan State University (42.716989, −84.462711; microbiota input for holoxenic condition). Holoxenic (Holo) plants were inoculated with soil slurry (10g soil/L of 0.5x LS), whereas axenic (Ax) plants were inoculated with 0.5x LS liquid medium.

### Genetic screen

Roughly 30,000 *Arabidopsis bak1-5 bkk1-1 cerk1-2* (*bbc*) seeds were mutagenized using 0.2% ethyl methanesulfonate (EMS). Mutagenized M1 seeds were sown on soil and allowed to grow to set seeds. Seeds from two to three M1 plants were pooled and approximately 1,700 pools were collected. This EMS population was estimated to cover the *Arabidopsis* genome more than 10 times. The primary screen was conducted by seedling flood-inoculation assay^57^. In brief, roughly fifty M2 seeds from each pool were sown onto 0.5x LS agar plates followed by flooding 3-week-old seedlings with 1×10^8^ CFU/mL *Pst* D28E (suspended in 0.25mM MgCl_2_ and 0.015% Silwet L-77) for 4 minutes. After 4 minutes, inoculum was removed, and plates were returned to Percival chambers for disease symptom development. Mutants showing signs of chlorosis and/or necrosis were transplanted to potting soil and transferred to a growth chamber to collect M3 seeds. Secondary screen was conducted by monitoring symptoms after either syringe-infiltration of *Pst* D28E to 4-week-old soil-grown M3 plants or after growth in holoxenic condition in the FlowPot gnotobiotic plant growth system^25^.

### Mutation identification using mapping-by-sequencing approach

To identify the causative mutation in the *grm1* mutant, a mapping-by-sequencing population was generated by backcrossing the *grm1* mutant with the *bbc* mutant. All four F1 plants showed *bbc*-like morphology suggesting that the mutant trait is recessive. F1 plants were selfed to produce F2 populations. Of 674 F2 plants screened, 633 had *bbc*-like morphology and 41 were *grm1*-like. This ratio deviates from the 3:1 single nuclear gene inheritance pattern. However, mapping-by-sequencing data does not support the idea that the mutant phenotype in *grm1* plants was caused by two or more unlinked loci as the allele frequency only peaks at around 7Mb on chromosome 5 (Extended Data Fig. 2a); the G to A mutation in the *TIP1* (*At5g20350*) gene is tightly associated with the *grm1* phenotype (Extended Data Fig. 2 and Extended Data Table 2). A literature search found that, interestingly, a similar genetic inheritance pattern deviation was observed in the characterization of the first mutant allele of *TIP1* (*tip1-1*^44^). The authors suggested that “deficiency in the number of homozygous *tip1* mutant seeds” as a potential cause of such inheritance ratio deviation.

### Genetic complementation of the *grm1* mutant

High fidelity PCR was performed using Col-0 genomic DNA as template and with a sense primer covering roughly 2kb upstream of the *At5g20350* start codon and an anti-sense primer roughly 1kb downstream of the stop codon. Cloned PCR product was inserted into pDONR207 entry vector and verified using Sanger sequencing. Recombination reaction was conducted using verified entry clone and pMDC123 destination vector to create the *TIP1* genomic clone driven by the *TIP1* native promoter. The construct was transformed into the *grm1* mutant plants via floral dipping method ^58^ using *Agrobacterium tumefaciens* GV3101 as the vehicle strain. See Extended Data Table 5 for the primer sequences used in this study.

### Quantification of endophytic leaf bacterial microbiota

Four-week-old potting soil-grown plants were sprayed with distilled water and fully covered with a clear dome to maintain high humidity (~95% RH) for 5 days (or 7 days for natural *Arabidopsis* accessions). After high humidity treatment, 1-3 leaves from each plant were surface sterilized with 4% bleach (0.33% active ingredient NaOCl) for one minute followed by two rinses with sterile water. Surface-sterilized leaves were blotted dry using paper towels and weighted. Sterile water was added to leaf samples, which were homogenized using TissueLyser II (QIAGEN) for 2 × 45 seconds at 30 Hz. Homogenized samples were serial-diluted and spotted onto R-2A plates (Sigma-Aldrich Cat. No. 17209) supplemented with cycloheximide (15 mg/L) and 0.5% MeOH to enumerate culturable colonies.

### Profiling microbial composition using 16S rDNA amplicon sequencing

For this experiment, all consumables and kits came from the same lot to avoid any background contamination or variations. Seeds of indicated *Arabidopsis* genotypes were surface sterilized using 15% bleach (1.2% active ingredient NaOCl) and washed twice using autoclaved water before sowing onto potting soil. Plants were grown in growth chambers. Four-week-old plants were sprayed with distilled water and kept at ≥95% relative humidity for five days. One to two leaves per plant were harvested, surface-sterilized using 4% bleach (0.33% active ingredient NaOCl) for one minute and followed by two washes using autoclaved Milli-Q water. Excess water on leaf surfaces was blotted dry, put into Safe-Lock Eppendorf tubes containing 3-mm zirconium beads (Glen Mills Inc), snap froze in liquid nitrogen and stored at −80 °C.

Total DNA (host and microbes) was extracted using DNeasy PowerSoil Pro Kit (Qiagen Cat. No. 47014) following the manufacturer’s protocol. Extracted DNA was used as template for PCR amplification of the v5/v6 region of 16S rRNA gene using 799F and 1193R primers (See Extended Data Table 5 for the sequence of primers) and high fidelity AccuPrime Taq DNA Polymerase (Invitrogen Cat. No. 12346086). Amplified products were run in 1% agarose gels to separate bacterial and chloroplast 16S rDNA amplicons (~400bp) from the mitochondrial 18S amplicon (~750bp). DNA in the ~400 bp band was recovered using the Zymoclean Gel DNA Recovery Kit (Zymo Research D4008). Concentration of recovered DNA was measured using the Quant-iT PicoGreen dsDNA Assay Kits (Invitrogen P7589) and normalized to 3-8 ng/μL for sample submission. Library preparation and sequencing using MiSeq platform (2 × 250 bp paired-end format) was conducted by the Genomic Core Facility at Michigan State University.

### 16S rDNA amplicon data processing

Raw Illumina data for 16S rDNA amplicon were processing as described previously^20^ using QIIME2 version 2022.2^59^. In brief, primer sequences were removed using Cutadapt^60^ followed by filtering, denoising and creating an ASV table by DADA2^61^. Taxonomic assignment of each ASV was performed using a Naïve Bayes classifier^62^ pre-trained on the SILVA 16S rRNA gene reference database^63,64^ (release 138) formatted for QIIME using RESCRIPt^65^. Unassigned sequences and sequences annotated as mitochondria and chloroplast were removed. Diversity analyses were performed within QIIME2. Samples were rarified to 1952 reads for calculating diversity metrics. The entire sequence analysis workflow is available on GitHub (https://github.com/BradCP/Roles-of-microbiota-in-autoimmunity-in-Arabidopsis).

### Bacterial infection assays

Four-week-old soil-grown plants were infiltrated with either *Pst* DC3000 at 1 to 2 × 10^5^ CFU/mL or *Pst* Δ*hrcC* at 2 to 3 × 10^6^ CFU/mL. After infiltration, plants were returned to growth chambers and kept at high humidity (~95% RH) for disease progression. Samples were harvested 3 days post inoculation (dpi) for *Pst* DC3000 or 5 dpi for *Pst* Δ*hrcC*. To determine bacterial population in leaves, leaf-discs were collected and ground in autoclaved Milli-Q water using a TissueLyser II (QIAGEN; 45 seconds at 30 Hz). Serial dilutions of the ground tissue were spotted onto low salt Luria Bertani (LB) plates (10 g/L Tryptone, 5 g/L Yeast Extract and 5 g/L NaCl; pH 7.0) with appropriate antibiotics. Colony forming units (CFUs) per cm^2^ were determined for each sample.

### RT-qPCR analysis gene expression

For gene expression analysis, plant tissues at the indicated conditions were harvested, snap frozen in liquid N_2_ and stored at −80°C until further processing.

Total RNA was extracted from plant tissues using TRIzol Reagent (Thermo Fisher Cat. No. 15596026) according to the manufacturer’s instructions. cDNA synthesis was accomplished in 10 µL volume with SuperScript IV VILO Master Mix (Thermo Fisher Cat. No. 11756050) according to the manufacturer’s instructions with 1µg total RNA as input. Upon reverse transcription, the product was diluted 5-fold using TE buffer (10 mM Tris-HCl pH 8.0, 1 mM EDTA). qPCR was performed in a 10 μL reaction volume containing 5 µL SYBR Green PCR master mix (Thermo Fisher Cat. No. 4309155), 500 nM of each primer, and 1 µL of template cDNA using a QuantStudio 3 real-tame PCR system (Applied Biosystems). *PP2AA3* was used for normalization. The primer sets used to quantify gene expression in this study are listed in Extended Data Table 5.

## Data availability

Raw Illumina data for 16S rDNA amplicon sequences for the *grm1* mutant and related controls are available in the Sequence Read Archive database (SRA) under BioProject PRJNA934331, accession numbers SAMN33271678 to SAMN33271728. Raw Illumina data for 16S rDNA amplicon sequences for Col-0, *tip1* and *snc1* are available in the SRA database under BioProject PRJNA934350, accession numbers SAMN33272493 to SAMN33272548.

## Code availability

There is no custom code generated for this study.

## Acknowledgements

We thank Dr. Xin Li for kindly providing us *snc1* seeds. We thank Michigan State University (MSU) Growth Chamber Facility, MSU Genomics Core Facility and Duke Phytotron for technical assistance. This work was supported by the Natural Sciences and Engineering Research Council of Canada (Y.T.C.) and United States National Institutes of Health (1R01AI155441; to S.Y.H.). S.Y.H. is an Investigator at Howard Hughes Medical Institute.

## Author information

### Contributions

Y.T.C. and S.Y.H. conceptualized and designed the project.

Y.T.C. led and conducted most of the experimental work.

C.A.T. generated EMS-mutagenized population of *Arabidopsis bbc* mutant.

C.A.T. and L.Z. carried out genetic screens and characterization of *grm* mutants.

B.C.P. contributed materials for gene expression analysis.

Y.T.C. and B.C.P. performed data analysis of the 16S rDNA amplicon sequencing.

Y.T.C and S.Y.H. wrote the manuscript with input from all authors.

## EXTENDED DATA FIGURES AND LEGENDS

**Extended Data Figure 1.**
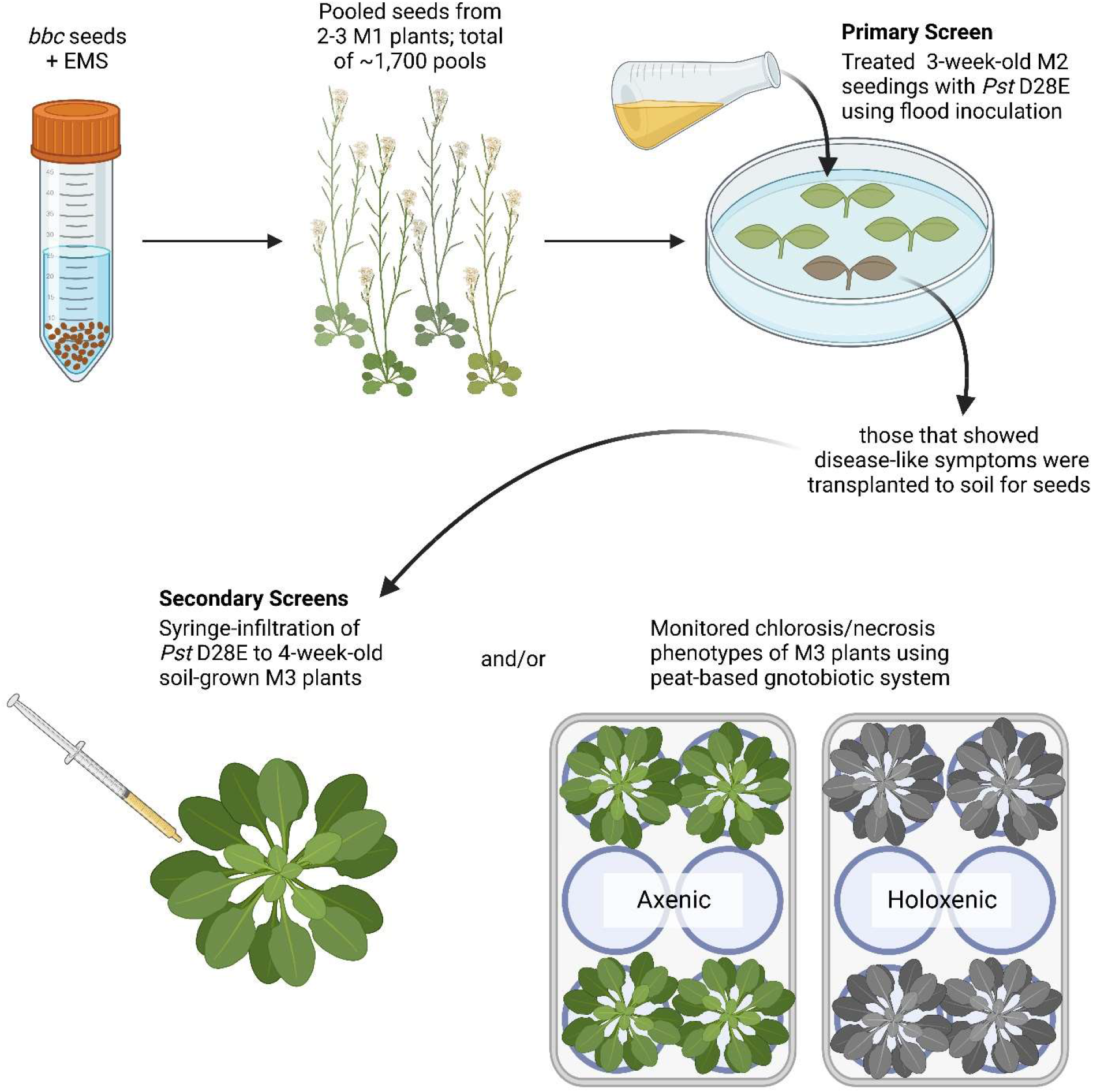
A schematic diagram of the genetic screen workflow. See Methods for detailed description of the genetic screen. Created with BioRender.com.

**Extended Data Figure 2.**
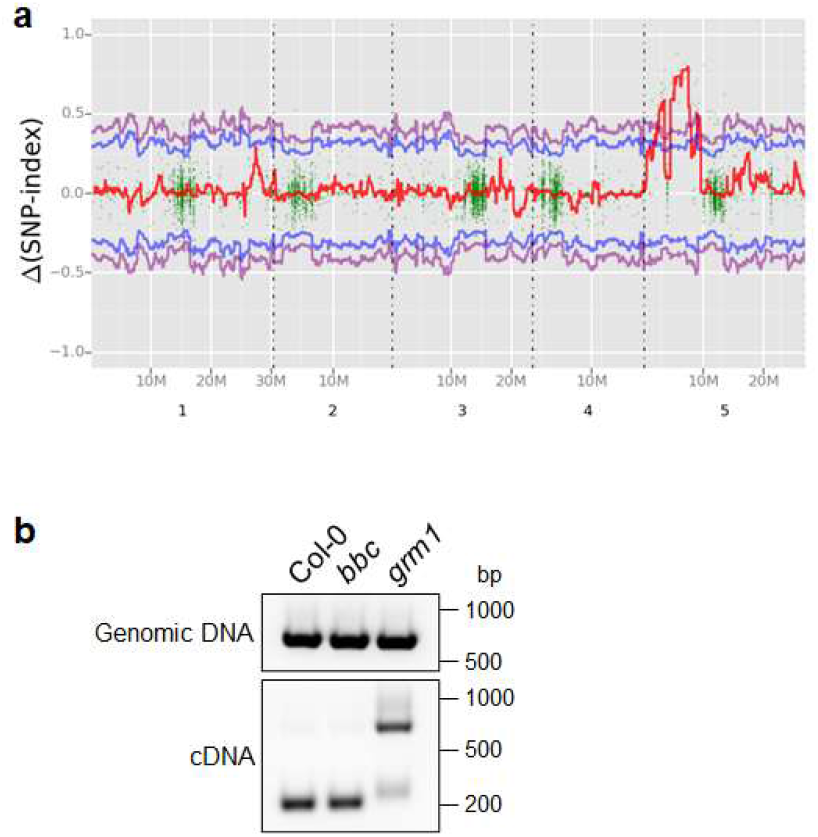
Mapping-by-sequencing to identify the causative mutation in the *grm1* mutant. **a,** *grm1* genomic mapping. Red line represents allele frequency. Blue and purple lines denote 95% and 99% confidence intervals, respectively. **b,** RT-PCR products using primers flanking the *grm1* mutation locus. Genomic (top panel) and complementary (bottom panel) DNA from indicated genotypes were used as templates.

**Extended Data Figure 3.**
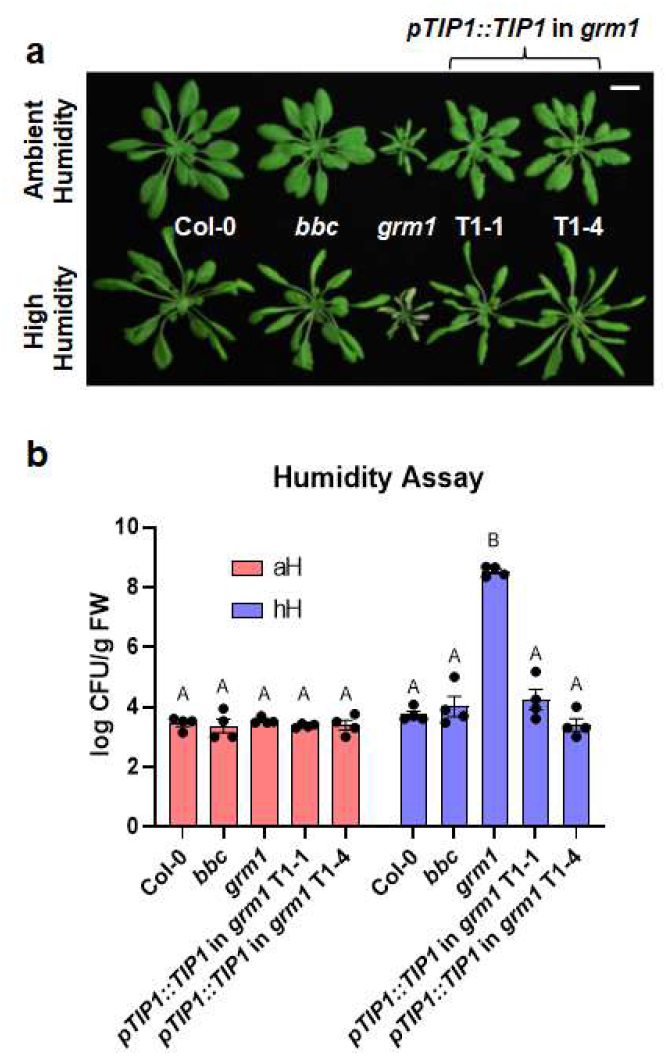
Transgene complementation of the *grm1* mutant. **a**, Images of four-week-old, potting soil-grown Col-0, *bbc*, *grm1* and two independent *grm1* complementation lines under ambient humidity (~50% RH basal control condition; upper panel) or high humidity (~95% RH; bottom panel) for five days. Scale bar equals 2 cm. **b,** Transgene complementation of the *grm1* mutant. Population sizes of leaf endophytic microbiota after five days of plant growth under ambient humidity (aH, ~50% RH basal control condition) or high humidity (~95% RH). Results represent the mean values ± SEM of four plants. Statistical analysis was performed using two-way ANOVA with Tukey’s HSD test. Experiment was independently performed twice with similar results.

**Extended Data Figure 4.**
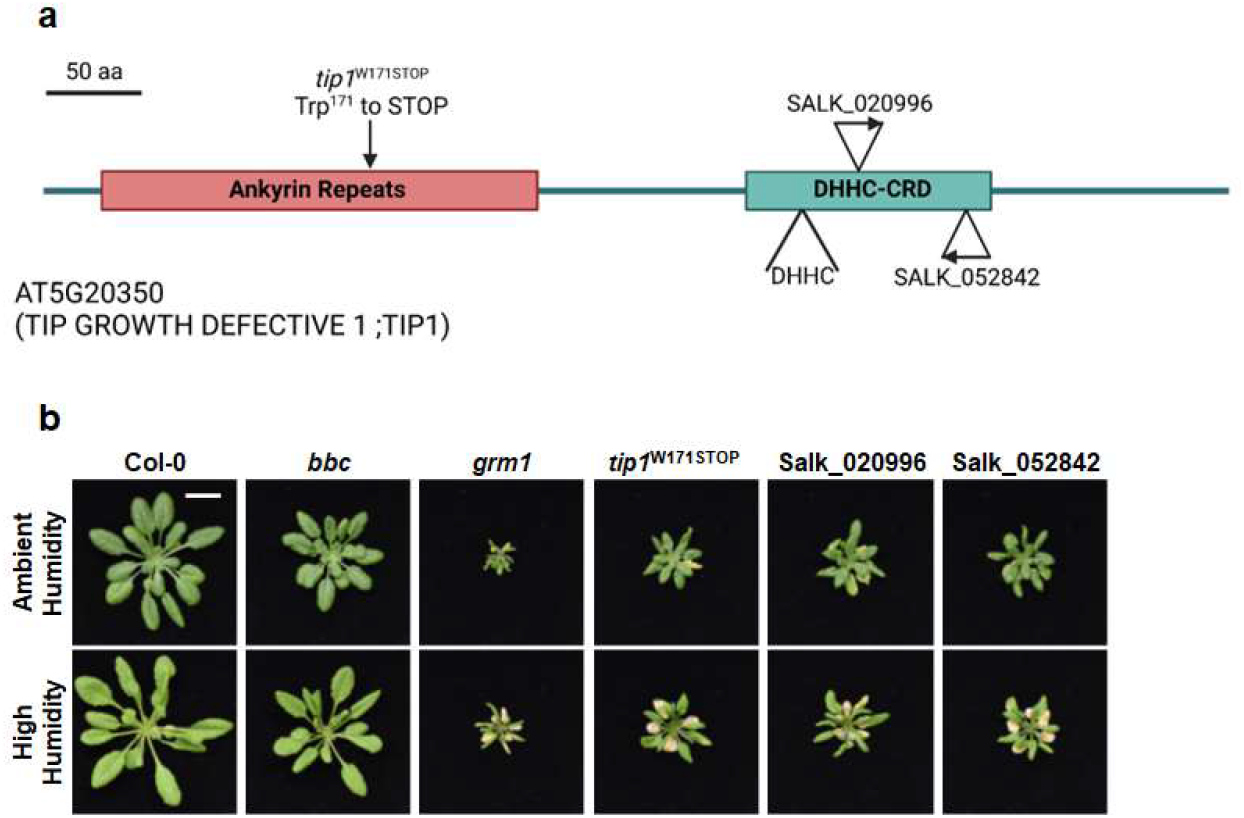
Appearance of *tip1* mutant alleles. **a,** A schematic diagram illustrating various mutant alleles of TIP1 protein. *tip1*^W171STOP^ is the allele isolated from this study which contains a G to A mutation at the splicing junction that is expected to cause a pre-mature STOP codon at amino acid residue Trp^171^ in the ankyrin-repeat domain. SALK_020996 and SALK_052842 have T-DNA insertions that would affect the DHHC cysteine-rich domain (DHHC-CRD). Created with BioRender.com. **b,** Images of four-week-old, potting soil-grown Col-0, *bbc*, *grm1* and various *tip1* single mutant plants under ambient humidity (~50% RH basal control condition; upper panel) or high humidity (~95% RH) for five days. Scale bar equals 2 cm.

**Extended Data Figure 5.**
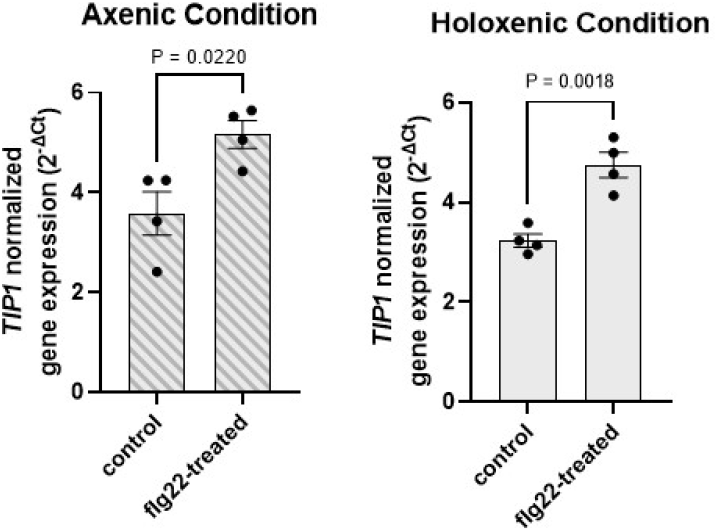
*TIP1* is induced by PTI elicitor flg22. Expression of *TIP1* after 90 minutes of 250 nM flg22 treatment. Plants were grown under axenic (left panel; with diagonal stripe pattern) or holoxenic (right panel) conditions in GnotoPots. Results represent the mean values ± SEM of four biological replicates. Statistical analysis was done by the Student’s *t*-test. Experiment was independently performed three times with similar results.

**Extended Data Figure 6.**
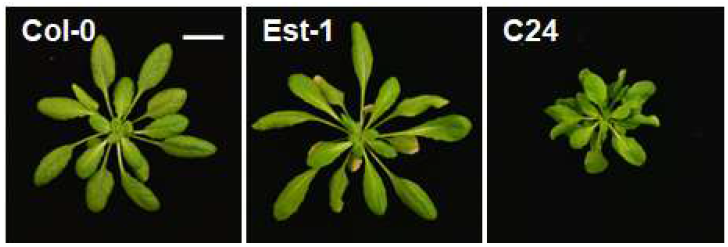
Appearance of 5-week-old, soil-grown Col-0, Est-1 and C24 plants.

**Extended Data Figure 7.**
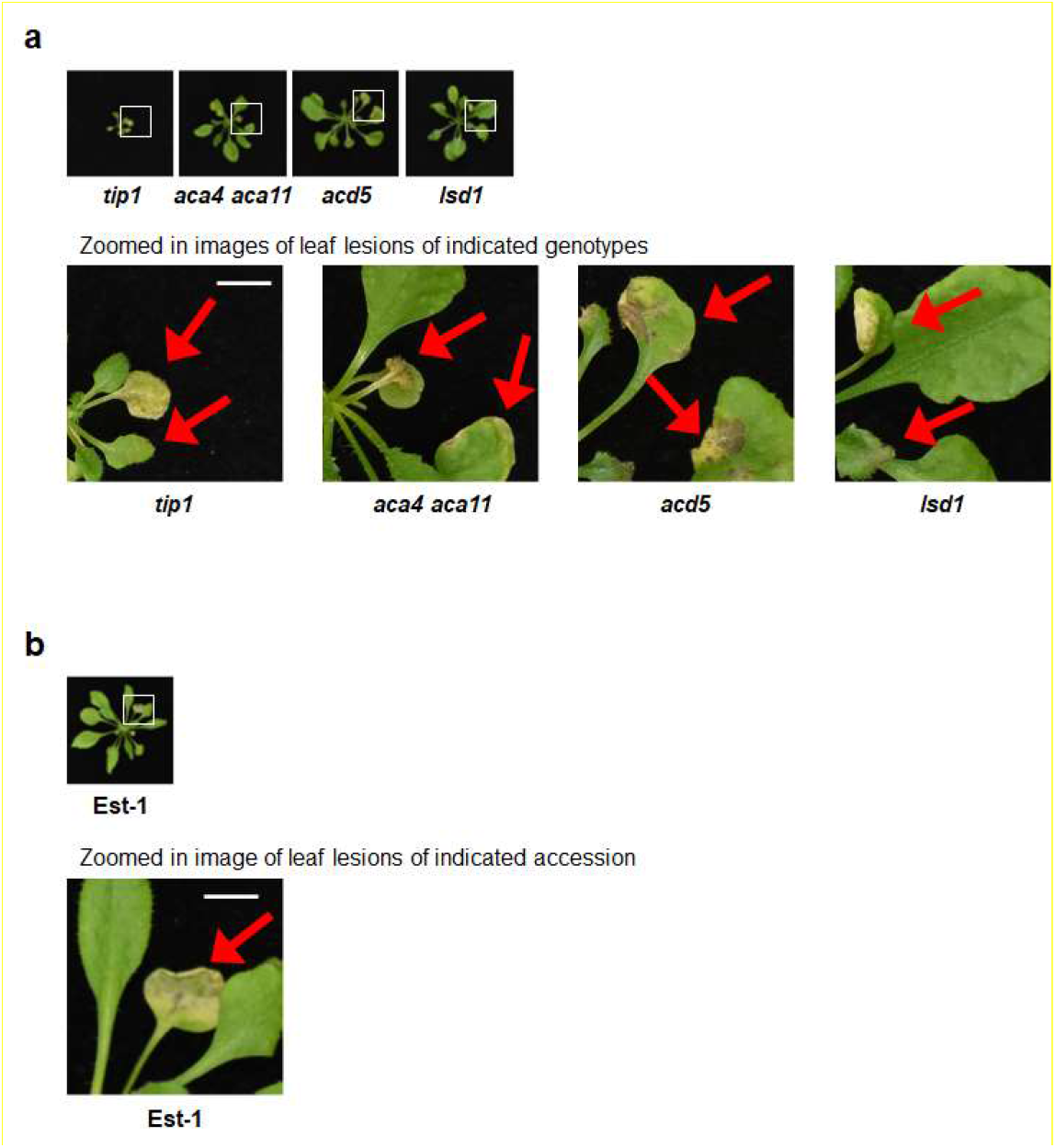
Zoomed in images of leaf lesions on plants grown in holoxenic conditions. **a,** Zoomed in images of indicated genotypes grown on GnotoPots under holoxenic conditions. Scale bar equals 0.5 cm. **b**, Zoomed in image of Est-1 grown on GnotoPots under holoxenic conditions. Scale bar equals 0.5 cm.

**Extended Data Figure 8.**
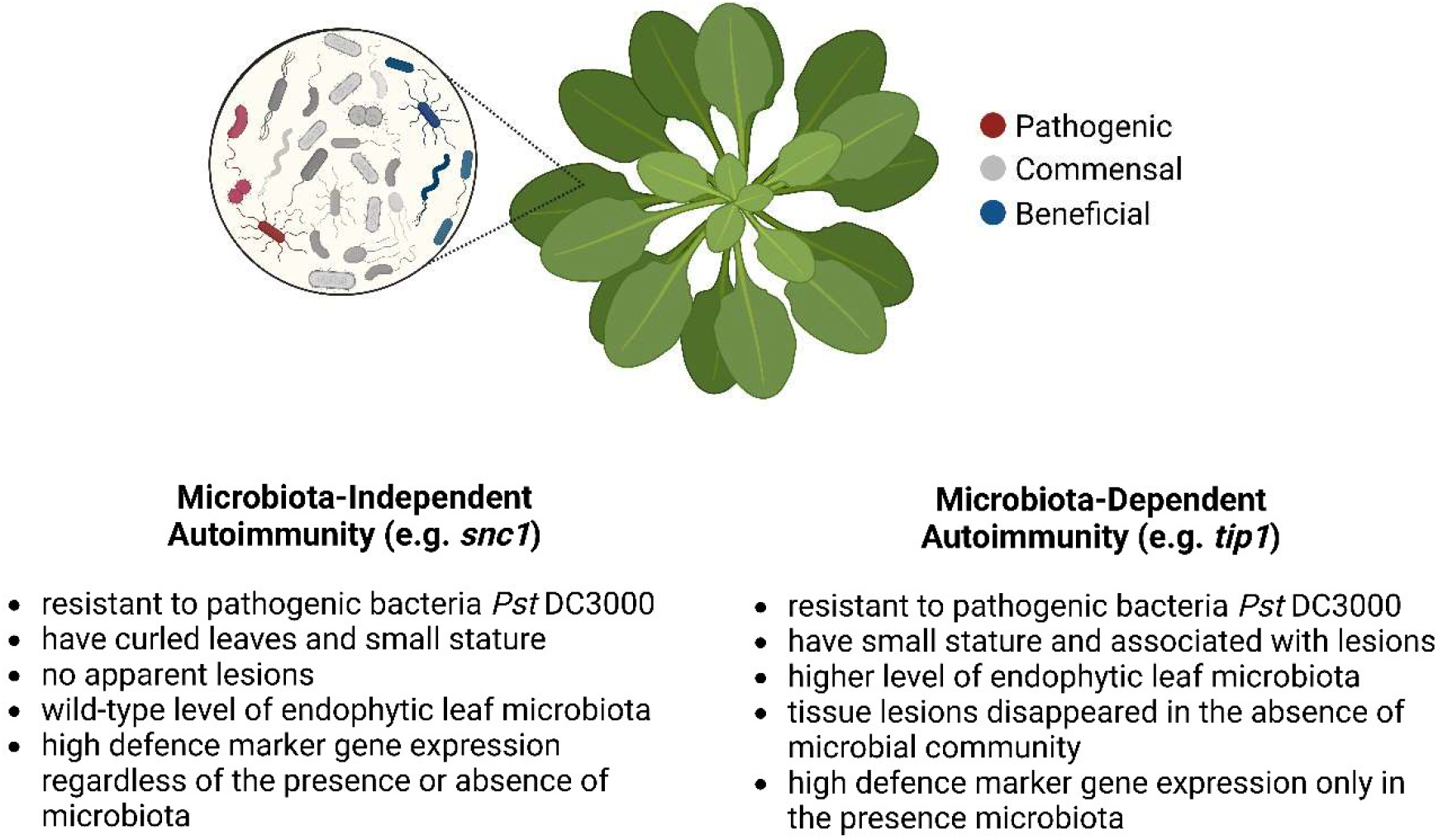
Characteristics of two types of autoimmunity in plants based on their microbiota dependency. Created with BioRender.com.

## Supplementary Information

**Extended Data Table 1. ASV counts of leaf endophytic microbiota in Col-0, *bbc* and *grm1* plants.**

**Extended Data Table 2. Mutations on chromosome 5 (from 4 to 12 Mbp) in the *grm1* mutant.**

**Extended Data Table 3. ASV counts of leaf endophytic microbiota in Col-0, *tip1* and *snc1* plants.**

**Extended Data Table 4. List of *Arabidopsis* mutants and accessions used in this study.**

**Extended Data Table 5. List of primers used in this study.**

## References

1 Staskawicz, B. J., Ausubel, F. M., Baker, B. J., Ellis, J. G. & Jones, J. D. Molecular genetics of plant disease resistance. Science 268, 661–667 (1995). https://doi.org:10.1126/science.7732374

2 Jones, J. D. & Dangl, J. L. The plant immune system. Nature 444, 323–329 (2006). https://doi.org:10.1038/nature05286

3 Ngou, B. P. M., Ding, P. & Jones, J. D. G. Thirty years of resistance: Zig-zag through the plant immune system. Plant Cell 34, 1447–1478 (2022). https://doi.org:10.1093/plcell/koac041

4 Lewis, J. D., Guttman, D. S. & Desveaux, D. The targeting of plant cellular systems by injected type III effector proteins. Semin Cell Dev Biol 20, 1055–1063 (2009). https://doi.org:10.1016/j.semcdb.2009.06.003

5 Dou, D. & Zhou, J. M. Phytopathogen effectors subverting host immunity: different foes, similar battleground. Cell Host Microbe 12, 484–495 (2012). https://doi.org:10.1016/j.chom.2012.09.003

6 Toruno, T. Y., Stergiopoulos, I. & Coaker, G. Plant-Pathogen Effectors: Cellular Probes Interfering with Plant Defenses in Spatial and Temporal Manners. Annu Rev Phytopathol 54, 419–441 (2016). https://doi.org:10.1146/annurev-phyto-080615-100204

7 Xin, X. F., Kvitko, B. & He, S. Y. Pseudomonas syringae: what it takes to be a pathogen. Nat Rev Microbiol 16, 316–328 (2018). https://doi.org:10.1038/nrmicro.2018.17

8 Zhou, J. M. & Zhang, Y. Plant Immunity: Danger Perception and Signaling. Cell 181, 978–989 (2020). https://doi.org:10.1016/j.cell.2020.04.028

9 Yuan, M., Ngou, B. P. M., Ding, P. & Xin, X. F. PTI-ETI crosstalk: an integrative view of plant immunity. Curr Opin Plant Biol 62, 102030 (2021). https://doi.org:10.1016/j.pbi.2021.102030

10 Huot, B., Yao, J., Montgomery, B. L. & He, S. Y. Growth-defense tradeoffs in plants: a balancing act to optimize fitness. Mol Plant 7, 1267–1287 (2014). https://doi.org:10.1093/mp/ssu049

11 He, Z., Webster, S. & He, S. Y. Growth-defense trade-offs in plants. Curr Biol 32, R634–R639 (2022). https://doi.org:10.1016/j.cub.2022.04.070

12 Muller, D. B., Vogel, C., Bai, Y. & Vorholt, J. A. The Plant Microbiota: Systems-Level Insights and Perspectives. Annu Rev Genet 50, 211–234 (2016). https://doi.org:10.1146/annurev-genet-120215-034952

13 Wang, N. R. & Haney, C. H. Harnessing the genetic potential of the plant microbiome. The Biochemist 42, 20–25 (2020).

14 Bulgarelli, D., Schlaeppi, K., Spaepen, S., Ver Loren van Themaat, E. & Schulze-Lefert, P. Structure and functions of the bacterial microbiota of plants. Annu Rev Plant Biol 64, 807–838 (2013). https://doi.org:10.1146/annurev-arplant-050312-120106

15 Paasch, B. C. & He, S. Y. Toward understanding microbiota homeostasis in the plant kingdom. PLoS Pathog 17, e1009472 (2021). https://doi.org:10.1371/journal.ppat.1009472

16 Hacquard, S., Spaepen, S., Garrido-Oter, R. & Schulze-Lefert, P. Interplay Between Innate Immunity and the Plant Microbiota. Annu Rev Phytopathol 55, 565–589 (2017). https://doi.org:10.1146/annurev-phyto-080516-035623

17 Nobori, T. et al. Dissecting the cotranscriptome landscape of plants and their microbiota. EMBO Rep, e55380 (2022). https://doi.org:10.15252/embr.202255380

18 Velasquez, A. C., Huguet-Tapia, J. C. & He, S. Y. Shared in planta population and transcriptomic features of nonpathogenic members of endophytic phyllosphere microbiota. Proc Natl Acad Sci U S A 119, e2114460119 (2022). https://doi.org:10.1073/pnas.2114460119

19 Xin, X. F. et al. Bacteria establish an aqueous living space in plants crucial for virulence. Nature 539, 524–529 (2016). https://doi.org:10.1038/nature20166

20 Chen, T. et al. A plant genetic network for preventing dysbiosis in the phyllosphere. Nature 580, 653–657 (2020). https://doi.org:10.1038/s41586-020-2185-0

21 Pfeilmeier, S. et al. The plant NADPH oxidase RBOHD is required for microbiota homeostasis in leaves. Nat Microbiol 6, 852–864 (2021). https://doi.org:10.1038/s41564-021-00929-5

22 Gimenez-Ibanez, S. et al. AvrPtoB targets the LysM receptor kinase CERK1 to promote bacterial virulence on plants. Curr Biol 19, 423–429 (2009). https://doi.org:10.1016/j.cub.2009.01.054

23 Schwessinger, B. et al. Phosphorylation-dependent differential regulation of plant growth, cell death, and innate immunity by the regulatory receptor-like kinase BAK1. PLoS Genet 7, e1002046 (2011). https://doi.org:10.1371/journal.pgen.1002046

24 Cunnac, S. et al. Genetic disassembly and combinatorial reassembly identify a minimal functional repertoire of type III effectors in Pseudomonas syringae. Proc Natl Acad Sci U S A 108, 2975–2980 (2011). https://doi.org:10.1073/pnas.1013031108

25 Kremer, J. M. et al. Peat-based gnotobiotic plant growth systems for Arabidopsis microbiome research. Nat Protoc 16, 2450–2470 (2021). https://doi.org:10.1038/s41596-021-00504-6

26 Alonso, J. M. et al. Genome-wide insertional mutagenesis of Arabidopsis thaliana. Science 301, 653–657 (2003). https://doi.org:10.1126/science.1086391

27 Gomez-Gomez, L., Felix, G. & Boller, T. A single locus determines sensitivity to bacterial flagellin in Arabidopsis thaliana. Plant J 18, 277–284 (1999). https://doi.org:10.1046/j.1365-313x.1999.00451.x

28 Boudsocq, M. et al. Differential innate immune signalling via Ca(2+) sensor protein kinases. Nature 464, 418–422 (2010). https://doi.org:10.1038/nature08794

29 Zhang, Y., Goritschnig, S., Dong, X. & Li, X. A gain-of-function mutation in a plant disease resistance gene leads to constitutive activation of downstream signal transduction pathways in suppressor of npr1-1, constitutive 1. Plant Cell 15, 2636–2646 (2003). https://doi.org:10.1105/tpc.015842

30 Cheng, Y. T. et al. Stability of plant immune-receptor resistance proteins is controlled by SKP1-CULLIN1-F-box (SCF)-mediated protein degradation. Proc Natl Acad Sci U S A 108, 14694–14699 (2011). https://doi.org:10.1073/pnas.1105685108

31 Wei, W. et al. The gene coding for the Hrp pilus structural protein is required for type III secretion of Hrp and Avr proteins in Pseudomonas syringae pv. tomato. Proc Natl Acad Sci U S A 97, 2247–2252 (2000). https://doi.org:10.1073/pnas.040570097

32 van Wersch, R., Li, X. & Zhang, Y. Mighty Dwarfs: Arabidopsis Autoimmune Mutants and Their Usages in Genetic Dissection of Plant Immunity. Front Plant Sci 7, 1717 (2016). https://doi.org:10.3389/fpls.2016.01717

33 Bruggeman, Q., Raynaud, C., Benhamed, M. & Delarue, M. To die or not to die? Lessons from lesion mimic mutants. Front Plant Sci 6, 24 (2015). https://doi.org:10.3389/fpls.2015.00024

34 Boursiac, Y. et al. Disruption of the vacuolar calcium-ATPases in Arabidopsis results in the activation of a salicylic acid-dependent programmed cell death pathway. Plant Physiol 154, 1158–1171 (2010). https://doi.org:10.1104/pp.110.159038

35 Liang, H. et al. Ceramides modulate programmed cell death in plants. Genes Dev 17, 2636–2641 (2003). https://doi.org:10.1101/gad.1140503

36 Zeng, H. Y. et al. The immune components ENHANCED DISEASE SUSCEPTIBILITY 1 and PHYTOALEXIN DEFICIENT 4 are required for cell death caused by overaccumulation of ceramides in Arabidopsis. Plant J 107, 1447–1465 (2021). https://doi.org:10.1111/tpj.15393

37 Dietrich, R. A., Richberg, M. H., Schmidt, R., Dean, C. & Dangl, J. L. A novel zinc finger protein is encoded by the Arabidopsis LSD1 gene and functions as a negative regulator of plant cell death. Cell 88, 685–694 (1997). https://doi.org:10.1016/s0092-8674(00)81911-x

38 Yang, H. et al. A mutant CHS3 protein with TIR-NB-LRR-LIM domains modulates growth, cell death and freezing tolerance in a temperature-dependent manner in Arabidopsis. Plant J 63, 283–296 (2010). https://doi.org:10.1111/j.1365-313X.2010.04241.x

39 Clough, S. J. et al. The Arabidopsis dnd1 “defense, no death” gene encodes a mutated cyclic nucleotide-gated ion channel. Proc Natl Acad Sci U S A 97, 9323–9328 (2000). https://doi.org:10.1073/pnas.150005697

40 Jurkowski, G. I. et al. Arabidopsis DND2, a second cyclic nucleotide-gated ion channel gene for which mutation causes the “defense, no death” phenotype. Mol Plant Microbe Interact 17, 511–520 (2004). https://doi.org:10.1094/MPMI.2004.17.5.511

41 Velasquez, A. C., Oney, M., Huot, B., Xu, S. & He, S. Y. Diverse mechanisms of resistance to Pseudomonas syringae in a thousand natural accessions of Arabidopsis thaliana. New Phytol 214, 1673–1687 (2017). https://doi.org:10.1111/nph.14517

42 Ton, J., Pieterse, C. M. & Van Loon, L. C. Identification of a Locus in Arabidopsis Controlling Both the Expression of Rhizobacteria-Mediated Induced Systemic Resistance (ISR) and Basal Resistance Against *Pseudomonas syringae* pv. *tomato*. Mol Plant Microbe Interact 12, 911–918 (1999). https://doi.org:10.1094/MPMI.1999.12.10.911

43 Todesco, M. et al. Natural allelic variation underlying a major fitness trade-off in Arabidopsis thaliana. Nature 465, 632–636 (2010). https://doi.org:10.1038/nature09083

44 Schiefelbein, J., Galway, M., Masucci, J. & Ford, S. Pollen tube and root-hair tip growth is disrupted in a mutant of Arabidopsis thaliana. Plant Physiol 103, 979–985 (1993). https://doi.org:10.1104/pp.103.3.979

45 Hemsley, P. A., Kemp, A. C. & Grierson, C. S. The TIP GROWTH DEFECTIVE1 S-acyl transferase regulates plant cell growth in Arabidopsis. Plant Cell 17, 2554–2563 (2005). https://doi.org:10.1105/tpc.105.031237

46 Singaraja, R. R. et al. HIP14, a novel ankyrin domain-containing protein, links huntingtin to intracellular trafficking and endocytosis. Hum Mol Genet 11, 2815–2828 (2002). https://doi.org:10.1093/hmg/11.23.2815

47 Babu, P., Deschenes, R. J. & Robinson, L. C. Akr1p-dependent palmitoylation of Yck2p yeast casein kinase 1 is necessary and sufficient for plasma membrane targeting. J Biol Chem 279, 27138–27147 (2004). https://doi.org:10.1074/jbc.M403071200

48 Kumar, M., Carr, P. & Turner, S. R. An atlas of Arabidopsis protein S-acylation reveals its widespread role in plant cell organization and function. Nat Plants 8, 670–681 (2022). https://doi.org:10.1038/s41477-022-01164-4

49 Hemsley, P. A., Weimar, T., Lilley, K. S., Dupree, P. & Grierson, C. S. A proteomic approach identifies many novel palmitoylated proteins in Arabidopsis. New Phytol 197, 805–814 (2013). https://doi.org:10.1111/nph.12077

50 Choi, S. C. et al. Gut microbiota dysbiosis and altered tryptophan catabolism contribute to autoimmunity in lupus-susceptible mice. Sci Transl Med 12 (2020). https://doi.org:10.1126/scitranslmed.aax2220

51 Cox, L. M. et al. Gut Microbiome in Progressive Multiple Sclerosis. Ann Neurol 89, 1195–1211 (2021). https://doi.org:10.1002/ana.26084

52 Chriswell, M. E. et al. Clonal IgA and IgG autoantibodies from individuals at risk for rheumatoid arthritis identify an arthritogenic strain of Subdoligranulum. Sci Transl Med 14, eabn5166 (2022). https://doi.org:10.1126/scitranslmed.abn5166

53 Yanai, A. et al. Palmitoylation of huntingtin by HIP14 is essential for its trafficking and function. Nat Neurosci 9, 824–831 (2006). https://doi.org:10.1038/nn1702

54 Lemonidis, K., Sanchez-Perez, M. C. & Chamberlain, L. H. Identification of a Novel Sequence Motif Recognized by the Ankyrin Repeat Domain of zDHHC17/13 S-Acyltransferases. J Biol Chem 290, 21939–21950 (2015). https://doi.org:10.1074/jbc.M115.657668

55 Lemonidis, K., MacLeod, R., Baillie, G. S. & Chamberlain, L. H. Peptide array-based screening reveals a large number of proteins interacting with the ankyrin-repeat domain of the zDHHC17 S-acyltransferase. J Biol Chem 292, 17190–17202 (2017). https://doi.org:10.1074/jbc.M117.799650

56 Verardi, R., Kim, J. S., Ghirlando, R. & Banerjee, A. Structural Basis for Substrate Recognition by the Ankyrin Repeat Domain of Human DHHC17 Palmitoyltransferase. Structure 25, 1337–1347 e1336 (2017). https://doi.org:10.1016/j.str.2017.06.018

57 Ishiga, Y., Ishiga, T., Uppalapati, S. R. & Mysore, K. S. Arabidopsis seedling flood-inoculation technique: a rapid and reliable assay for studying plant-bacterial interactions. Plant Methods 7, 32 (2011). https://doi.org:10.1186/1746-4811-7-32

58 Clough, S. J. & Bent, A. F. Floral dip: a simplified method for Agrobacterium-mediated transformation of Arabidopsis thaliana. Plant J 16, 735–743 (1998). https://doi.org:10.1046/j.1365-313x.1998.00343.x

59 Bolyen, E. et al. Reproducible, interactive, scalable and extensible microbiome data science using QIIME 2. Nat Biotechnol 37, 852–857 (2019). https://doi.org:10.1038/s41587-019-0209-9

60 Martin, M. Cutadapt removes adapter sequences from high-throughput sequencing reads. *EMBnet*. journal 17, 10–12 (2011).

61 Callahan, B. J. et al. DADA2: High-resolution sample inference from Illumina amplicon data. Nat Methods 13, 581–583 (2016). https://doi.org:10.1038/nmeth.3869

62 Mohammad, N. S., Nazli, R., Zafar, H. & Fatima, S. Effects of lipid based Multiple Micronutrients Supplement on the birth outcome of underweight pre-eclamptic women: A randomized clinical trial. Pak J Med Sci 38, 219–226 (2022). https://doi.org:10.12669/pjms.38.1.4396

63 Yilmaz, P. et al. The SILVA and “All-species Living Tree Project (LTP)” taxonomic frameworks. Nucleic Acids Res 42, D643–648 (2014). https://doi.org:10.1093/nar/gkt1209

64 Quast, C. et al. The SILVA ribosomal RNA gene database project: improved data processing and web-based tools. Nucleic Acids Res 41, D590–596 (2013). https://doi.org:10.1093/nar/gks1219

65 Robeson, M. S., 2nd et al. RESCRIPt: Reproducible sequence taxonomy reference database management. PLoS Comput Biol 17, e1009581 (2021). https://doi.org:10.1371/journal.pcbi.1009581

